# *In Silico* Evaluation of Cell Therapy in Acute versus Chronic Infarction: Role of Automaticity, Heterogeneity and Purkinje in Human

**DOI:** 10.1101/2023.12.11.570780

**Authors:** Leto Luana Riebel, Zhinuo Jenny Wang, Hector Martinez-Navarro, Cristian Trovato, Julia Camps, Lucas Arantes Berg, Xin Zhou, Ruben Doste, Rafael Sachetto Oliveira, Rodrigo Weber dos Santos, Jacopo Biasetti, Blanca Rodriguez

## Abstract

Human-based modelling and simulation offer an ideal testbed for novel medical therapies to guide experimental and clinical studies. Myocardial infarction (MI) is a common cause of heart failure and mortality, for which novel therapies are urgently needed. Although cell therapy offers promise, electrophysiological heterogeneity raises pro-arrhythmic safety concerns, where underlying complex spatio-temporal dynamics cannot be investigated experimentally. After demonstrating credibility of the modelling and simulation framework, we investigate cell therapy in acute versus chronic MI, and the role of cell heterogeneity, scar size and the Purkinje system.

Simulations agreed with experimental and clinical recordings from ionic to ECG dynamics in acute and chronic infarction. Following cell delivery, spontaneous beats were facilitated by heterogeneity in cell populations, chronic MI due to tissue depolarisation, and slow sinus rhythm. Subsequent re-entrant arrhythmias occurred, in some instances with Purkinje involvement, and their susceptibility was enhanced by impaired Purkinje-myocardium coupling, large scars, and acute infarction.

We conclude that homogeneity in injected cell populations minimises their spontaneous beating, which is enhanced by chronic MI, whereas a healthy Purkinje-myocardium coupling is key to prevent subsequent re-entrant arrhythmias, particularly for large scars.

## Introduction

Heart failure, commonly the result of myocardial infarction (MI), remains a leading cause of mortality worldwide with novel treatments needed to reduce the clinical burden. During MI, an insufficient blood and nutrient supply causes the affected tissue and bordering regions to undergo electrophysiological as well as structural remodelling. As a result, the heart’s contractile force decreases and adverse remodelling occurs also in the remote myocardium (Holmes et al., 2005; Sutton & Sharpe, 2000). Although regenerative cell therapy has been shown as a promising treatment in pre-clinical studies (Foo et al., 2018; Querdel et al., 2021; Zimmermann et al., 2006), safety concerns remain over the cells’ immaturities facilitating ventricular arrhythmias, as observed experimentally (Chong et al., 2014; Romagnuolo et al., 2019). While animal models present a valuable tool to evaluate new therapies, they are costly, time-consuming, exhibit limited control over experimental conditions and raise ethical and translational concerns. This limits progress in optimising therapies for MI.

Human modelling and simulation of cardiac electrophysiology is a mature and well-established technology enabling multiscale investigations into disease mechanisms (Arevalo et al., 2013; Martinez-Navarro et al., 2019; Wang et al., 2021), therapeutic interventions such as ablation (Roney et al., 2020), diagnostics (O’Hara et al., 2022) and pharmacological treatment (Dasí et al., 2022). While several studies have provided insights into specific pro-arrhythmic mechanisms following cell delivery (Fassina et al., 2022, 2023; Gibbs et al., 2023; Yu et al., 2019, 2021), they so far did not include thorough multi-scale validation from ionic to ECG level, nor did they address crucial factors such as disease progression, scar size and cell heterogeneity caused by differences in differentiation and purification protocols (Ban et al., 2017; Jiang et al., 2020).

The goal of this study is two-fold: 1) to develop and establish the credibility of a state-of-the-art human modelling and simulation framework for personalised electrophysiological investigations including the Purkinje system, through multiscale validation with experimental and clinical data; 2) to evaluate key modulators of spontaneous beating and re-entrant arrhythmias following cell delivery in three infarct stages (acute, healing and chronic), considering variability in cell heterogeneity, scar size and the role of the Purkinje system.

## Results

### Credibility of human modelling and simulation from ionic to ECG dynamics

The anatomically accurate human biventricular electrophysiology models post-MI incorporate state-of-the-art human cellular models of Purkinje and ventricular electrophysiology, with credibility for healthy and MI conditions supported by data in supplementary table S1 and previous studies (Tomek et al., 2019; Trovato et al., 2020; Wang et al., 2021; Zhou et al., 2022). Simulated ECGs in sinus rhythm following activation of the Purkinje system are in agreement with clinical recordings across MI progression (Bousseljot et al., 2009; Chew et al., 2018; Nable & Brady, 2009), as illustrated in Figure 1 and supplementary Figures S1 and S2. Both simulated and clinical ECGs in acute and healing MI exhibited fractured QRS-complexes and inverted T-waves (Figure 1A&B). Chronic MI ECGs exhibited increased QRS-complex duration, persistent ST-segment elevation and normalised T-waves (Figure 1 and supplementary Figure S3C).

ECGs before and after cell delivery during acute and healing MI were almost identical, as expected due to similar conduction velocity (CV) and repolarisation times of human stem cell-derived cardiomyocytes (hPSC-CMs) and the infarct (Figure 1C). In chronic MI, hPSC-CM delivery caused a small increase in ST-segment elevation in leads V1 to V3 and T-wave depression in leads V4 and V5. This was caused by hPSC-CMs improving conduction in the electrically non-active infarct. hPSC-CM population heterogeneity had no effect on ECG signals under sinus rhythm across all scar sizes and infarct stages (see supplementary Figure S4).

**Figure 1.**
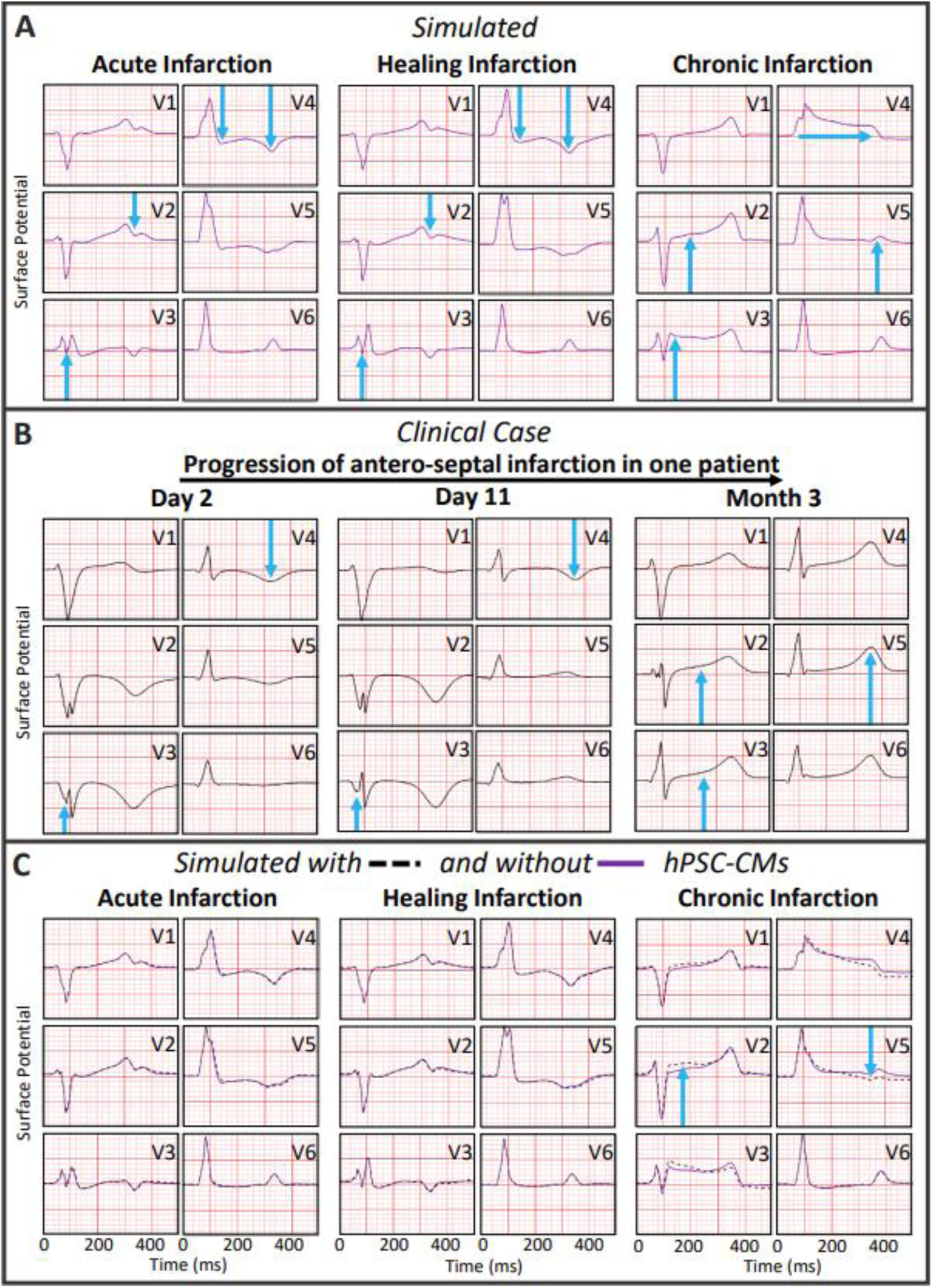
Comparison of simulated (A) and clinical ECGs (B) for acute, healing and chronic infarction, and following cell therapy (C). Clinical ECGs from the PTB Diagnostic ECG Database, https://physionet.org/content/ptbdb/1.0.0/ (Bousseljot et al., 2009; Goldberger et al., 2000), of patient 033 1, 10 and 89 days after anteroseptal infarction. Blue arrows highlight key changes compared to the healthy control. In C), simulated ECGs before (purple solid lines) and after (black dotted lines) cell delivery are compared. Shown simulations are in the medium scar and high injected cell heterogeneity with 50% ventricular-like, 25% atrial and 25% nodal-like phenotypes.

### Spontaneous beating is favoured by heterogeneous injected cell populations, depolarised infarcts and slow sinus beating

Figure 2 reports spontaneous activity in homogeneous versus heterogeneous stem cell populations injected in acute, healing and chronic infarctions at three scar sizes. Heterogeneous hPSC-CM populations with atrial and nodal-like phenotypes favoured ectopic activity, whereas homogeneous ventricular-like hPSC-CMs populations did not beat spontaneously for any scenario (see Figure 2A). Spontaneous hPSC-CM activity arose from infarct centres, where injections were simulated (Figure 2B and supplementary Figure S5).

The moderately heterogeneous cell populations (with 80% ventricular-like hPSC-CMs) beat spontaneously only in chronic MI, and its rate was the fastest for the medium scar, to become considerably slower for the larger scar (Figure 2A, middle column). Chronic MI promoted spontaneous beating through depolarisation of hPSC-CM diastolic membrane potential by electrotonic coupling with the infarct. Hence, an increase in scar size from small to medium led to faster spontaneous rates due to more depolarised membrane potentials (Figure 2A and supplementary Figure S6). However, being electrically inactive, large scars in chronic MI delayed the propagation of the spontaneous beat (Figure 2B, low action potential amplitude in the infarct in chronic MI compared to acute MI). This resulted in large current sinks dampening the spontaneous activity, as highlighted by a reduced upstroke velocity of hPSC-CMs action potentials for larger scars (1.2, 0.8, and 0.4 mV/ms in the small, medium, and large chronic scar for the moderately heterogeneous population).

The most heterogeneous population (50% ventricular-like hPSC-CMs) beat spontaneously in all post-MI scenarios (Figure 2A, last column and supplementary video SV1), and increasing scar size had very small effects on the beating times (+/-10ms, Figure 2A, last column). These much smaller differences in beating times can be explained by a faster spontaneous upstroke velocity (2.9, 2.6, and 2.2 mV/ms in the small, medium, and large chronic scar), highlighting the reduced current sink effects of the surrounding tissue. This increase in upstroke velocity with higher heterogeneity may appear counter intuitive as, in single cell, the ventricular-like phenotype had the fastest upstroke velocity (20.6 versus 8.9 and 4.3 mV/ms in the atrial and nodal-like phenotypes). However, in tissue, the quicker beating of atrial and nodal-like phenotypes drove the spontaneous depolarisation, having to excite not only the native ventricular tissue but also the slower beating ventricular-like hPSC-CMs. Hence, our results imply that heterogeneity in the injected cell population not only increases the rate of the spontaneous beating, but also its strength.

Furthermore, as illustrated in Figure 2C, spontaneous beating slowed down with increasing heart rate (from 1950 to 2250 ms for 50 to 100 bpm for the ventricular-like phenotype). Mechanistic investigation showed that this is caused by a depletion of the SR calcium deposit (Figure 2C, left, and more details in supplementary Figure S7). Accordingly, the peak of SR calcium release during the following spontaneous beat decreased from 0.15 to 0.08 mM/s. Faster spontaneous beats after slower heart rates were also observed in the biventricular simulations, with the medium scar during the acute stage shown as an example in the right panel (green circles) of Figure 2C.

**Figure 2.**
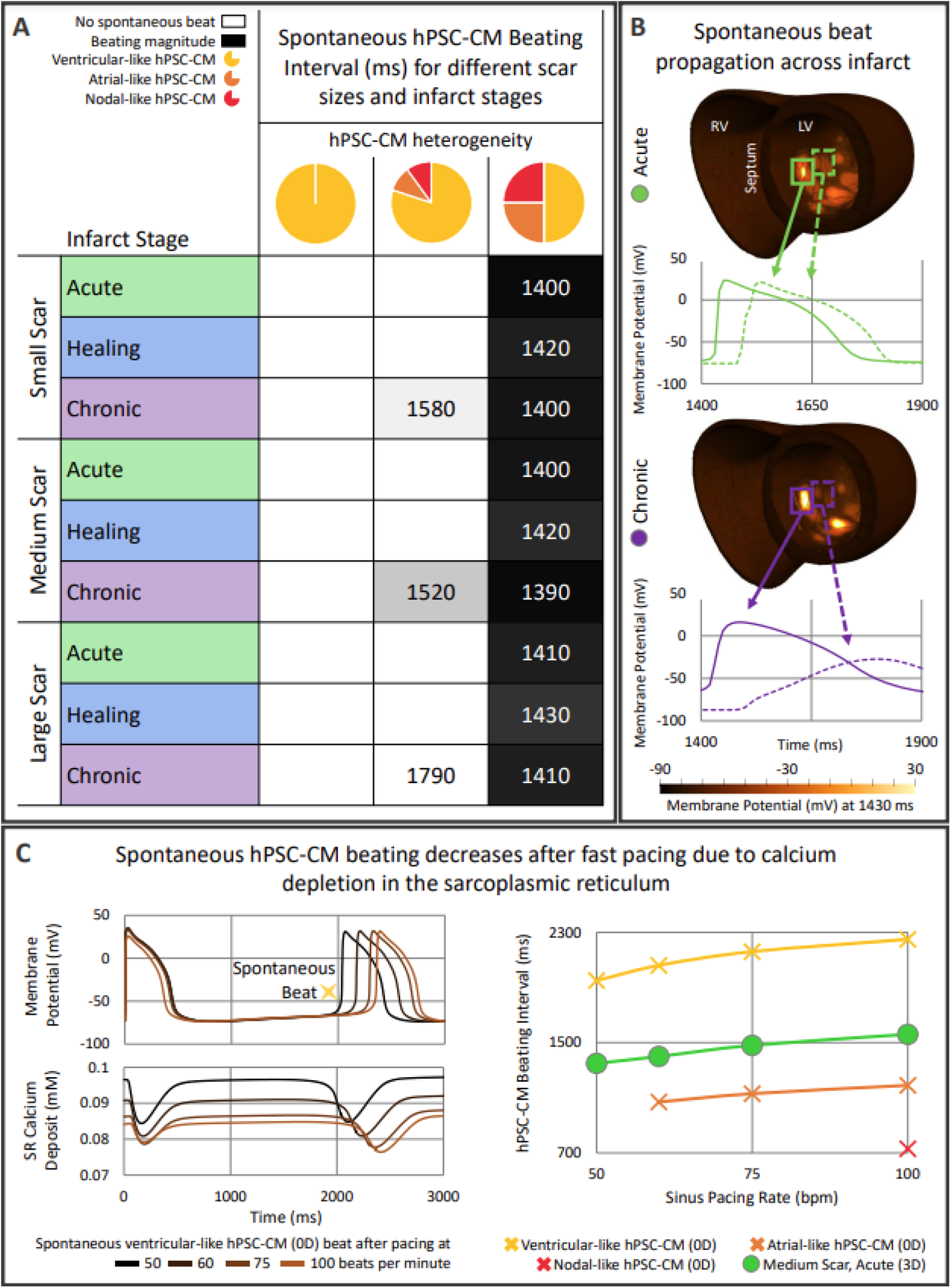
Modulators of spontaneous hPSC-CM beats. A) Spontaneous hPSC-CM beating intervals after the last sinus beat for increasing hPSC-CM population heterogeneity across infarct stages and sizes; dark shades indicate faster beating. B) Action potentials of infarcted cells from dense (solid lines) and sparse (dotted lines) hPSC-CM areas with the most heterogeneous hPSC-CM population in medium-sized acute (green) and chronic (purple) MI; left (LV) and right (RV) ventricle indicated. C) Effect of sinus heart rate on spontaneous hPSC-CM beats. Left: Single cell action potential and sarcoplasmic reticulum calcium deposit of ventricular-like hPSC-CM. Right: hPSC-CM beating intervals in single cell ventricular, atrial and nodal-like hPSC-CM phenotypes (0D) and coupled in the medium scar biventricular model with the most heterogeneous population in acute MI (3D). Spontaneous single cell atrial and nodal-like hPSC-CMs beating exceeded pacing below 60 and 100 bpm, respectively.

### Impaired Purkinje propagation increases arrhythmia susceptibility following spontaneous hPSC-CM beats

Re-entrant arrhythmia formation after spontaneous hPSC-CM activity under bradycardic heart rates was explored in conjunction with healthy versus impaired retrograde Purkinje propagation (Figure 3). Spontaneous hPSC-CM beating did not lead to the establishment of re-entry since the native tissue was already completely repolarised as shown in Figure 3A, and propagation engulfed the ventricles without the occurrence of unidirectional block (the first condition for re-entrant arrhythmias). However, spontaneous beating altered repolarisation patterns and provided the substrate for re-entrant arrhythmias following bradycardic beats through the bundle of His (between 250 to 450 ms after the spontaneous beat, Figure 3B). As shown in Figure 3C, the bradycardic beat met conduction block in the infarcted area still refractory from the spontaneous beat. For healthy Purkinje propagation, two re-entrant arrhythmias occurred in the large scar during the acute and chronic stage (Figure 3B), enabled by the wavefront retrogradely propagating through a Purkinje-myocyte junction (PMJ). In both cases, this PMJ was located within one of the scar cores, once in the left ventricular septo-apical wall and once on the antero-septal wall (see supplementary video SV2). The re-entry observed in the large scar during the acute stage was sustained for 3000 ms, with wavefronts re-entering also through the antero-basal and through the epicardial antero-apical wall (see left column Figure 3C).

Impaired retrograde Purkinje propagation (Figure 3, right column) promoted re-entry induction as shown by a widened vulnerable window for re-entry, which combining all infarct stages and scar sizes was 900 ms versus 20 ms with Purkinje retrograde propagation. This was because spontaneous beats could not propagate through the Purkinje system to remote regions, thereby increasing activation and repolarisation heterogeneity, the substrate for re-entry establishment. The vulnerable window increased with larger scar size (130 ms versus 420 ms for small versus large scar). Overall, acute MI was the most pro-arrhythmic, accounting for 54, 54 and 48 % of re-entrant arrhythmias in the small, medium and large scar, respectively. The chronic stage was the least pro-arrhythmic in all scar sizes, accounting for 0, 6 and 23 % of re-entries in the small, medium and large scar, respectively. The proportion of sustained arrhythmias continuing for at least two consecutive re-entry cycles increased from 31 over 69 to 81 % in the small, medium and large scar, respectively (Figure 3B). Re-entries were sustained for over 5000 ms, as shown in the ECG in the right column of Figure 3C. We identified different re-entrant circuit locations in the septum, anterior wall and apex, which varied across the different scar sizes, as illustrated in Figure 3C left and right column. No preference of re-entrant circuit location for any infarct size or stage was apparent. However, re-entry complexity increased with increasing scar size (see supplementary Figure S8). In the small scar, we observed at most two re-entrant circuit locations per re-entrant simulation. In the medium and large scar, 25 % of re-entries occurred through three or more sites simultaneously. The number of re-entrant circuit sites was similar during the acute and healing stages across all scar sizes. Re-entries during the chronic stage showed the least complexity; for example, in the large scar most re-entries occurred through a single circuit in the septal wall.

To summarise, while healthy Purkinje propagation enabled the spontaneous hPSC-CM beat to activate remote regions quickly and uniformly, impaired retrograde propagation caused a heterogeneous refractory substrate, which can be seen when comparing the left and right column of Figure 3A at 1500 ms. Furthermore, when retrogradely activated, the increased refractoriness of Purkinje cells compared to the ventricular tissue prevented re-excitation of the bundle of His for bradycardic intervals below 1780 ms when most of the ventricles had already repolarised, reducing the substrate for conduction block and re-entry. Overall, our results show how impaired retrograde Purkinje propagation can cause dyssynchronous propagation of the spontaneous beat and accordingly larger repolarisation gradients in the ventricles. In our study, these resulted in a multitude of re-entrant pathways and an overall increased re-entry inducibility, sustainability and complexity.

**Figure 3.**
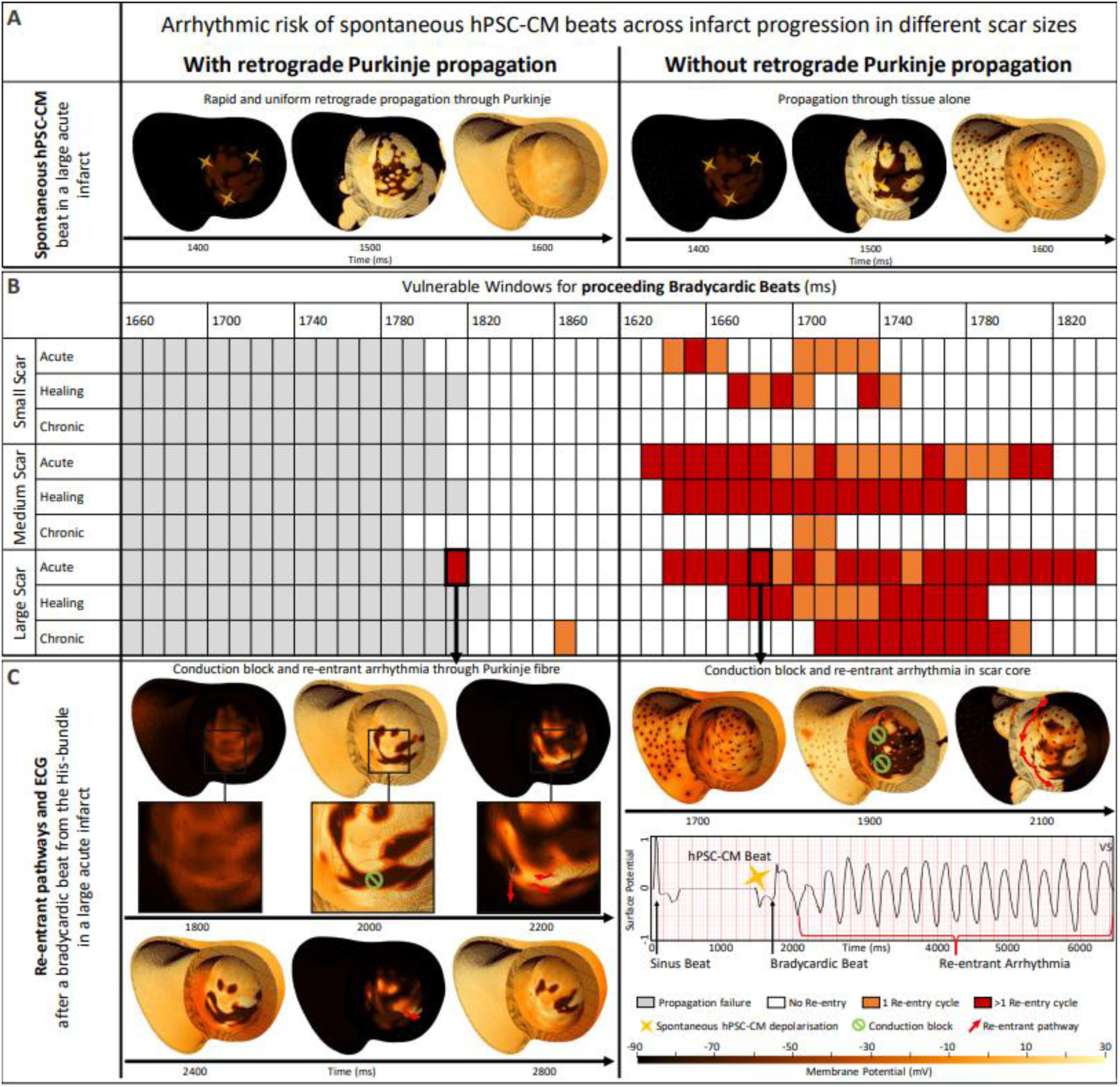
Depolarisation of the ventricles through spontaneous hPSC-CM beats and subsequent arrhythmia inducibility and sustainability. A) Emergence of spontaneous hPSC-CM beats in the large scar during the acute stage with (left) and without (right) retrograde Purkinje propagation. B) Vulnerable windows for the small, medium and large scar across the acute, healing and chronic stages with and without retrograde propagation through the Purkinje network. C) Propagation of electrical excitation following a proceeding bradycardic beat in the large scar during the acute stage post-MI with and without retrograde Purkinje propagation. The dark spots in the scenarios with retrograde Purkinje propagation disabled indicate the PMJs, showing that current was not able to flow from the myocardial tissue into the PMJs.

## Discussion

The first contribution of this study is the human multiscale modelling and simulation framework to investigate post-MI mechanisms and treatments, such as regenerative cell therapy. The credibility of the framework is supported by extensive consistency of simulation results in the present and previous studies (Zhou et al., 2021, 2022) with experimental and clinical data from ionic, cellular and importantly ECG dynamics from acute to chronic MI. The precise control in modelling and simulation studies allowed us to break down the key factors and mechanisms modulating the therapy’s safety. Specifically, here we investigated the interplay between infarct stage and size, Purkinje coupling and immature stem cell heterogeneity in modulating the pro-arrhythmic substrate following cell therapy.

Our key findings are as follows. Firstly, homogeneity in the injected cell population is crucial to avoid spontaneous beating activity, which is enhanced by depolarised infarcts such as in chronic MI, and slow beating rates. Secondly, subsequent arrhythmic burden due to re-entrant arrhythmias was highest in acute MI with large scars, and with impaired retrograde Purkinje coupling. Taken together, our simulations suggest that ventricular-like only cell populations in healing scars are less likely to produce pro-arrhythmic spontaneous beats, and that healthy Purkinje-myocardium coupling is also crucial.

Our simulations indicate that a substantial proportion of atrial and nodal-like phenotypes (>20%) is necessary for spontaneous beats to emerge in the ventricles. Heterogeneity within hPSC-CM cultures has been shown experimentally resulting from imperfect purification protocols and depending on maturation and differentiation protocols (Ban et al., 2017; Jiang et al., 2020). Heterogeneities include the action potential duration (APD), the diastolic membrane potential (DMP) and the spontaneous beating interval, and as shown in (Doss et al., 2012; He et al., 2003; Ma et al., 2011), may be even larger than investigated here. Such heterogeneities may have crucial arrhythmic implications which need to be assessed before therapy implementation, especially as longitudinal data of the cells after delivery are scarce.

A previous *in silico* study showed that the risk of spontaneous hPSC-CM beats increases with higher hPSC-CM density, larger hPSC-CM clustering and injection proximal rather than distal to the infarct (Yu et al., 2019). In our study, we modelled the spatial distribution of hPSC-CM based on experimental observations from (Poch et al., 2022), rather than varying density, clustering and injection locations. Therefore, although the number of delivered cells relative to the scar size was maintained in our study, absolute cell number, density and clustering varied between the different scar geometries. For example, our medium scar contained a scar core at the antero-septal wall towards the apex where the heart wall is very thick causing more cells to cluster there. According to Yu et al., this might influence the spontaneous beating rate, but as shown in our results, the spontaneous beats emerged from similar sites across all scars, suggesting this effect to be secondary here.

Although cell delivery is aimed at patients with chronic infarction, exploring the safety of cell injection across earlier infarct stages is crucial, since delivery protocols remain to be optimised and re-infarction cannot be ruled out. In our study, chronic infarction, modelled as conductive but non-excitable, promoted spontaneous beating but large scars slowed its upstroke and propagation. According to Gibbs et al. slow-conducting passive scars may reduce the occurrence of spontaneous beats compared to non-conducting scars (Gibbs et al., 2023). Results by Fassina et al., suggest that cells with a CV 20 % or less that of healthy myocardium, like chosen here, may promote arrhythmic events (Fassina et al., 2022, 2023).

While some experimental studies have suggested the stem cells’ rapid automaticity to directly facilitate ventricular tachycardia (Marchiano et al., 2023), our study showed how the spontaneous beats may facilitating re-entrant arrhythmias, also involving the Purkinje system. Our simulations confirm that large scars may be more pro-arrhythmic than small scars and further suggest that the increased repolarisation time of the Purkinje network and its ability to uniformly distribute the spontaneous beat across the rest of the ventricles, may be protective against some arrhythmias. Increased repolarisation times in Purkinje cells compared to ventricular cardiomyocytes have been shown experimentally (Boyden et al., 2010), replicated in the single cell models used here. Rapid Purkinje conduction eliminated some re-entrant pathways while enabling new ones, which has been highlighted in other *in silico* studies (Deo et al., 2009; Jian et al., 2022). As in our study we only examined completely enabled or completely disabled retrograde Purkinje propagation, future studies may investigate impaired conductivity in specific PMJs relative to infarcted areas. Overall, our investigation of the Purkinje network highlights its importance as a key modulator of arrhythmic risk, so far rarely included in computational studies.

## Methods

### Human multiscale modelling and simulation framework

A human multiscale modelling and simulation framework (shown in Figure 4) was developed and evaluated using experimental and clinical data from ionic to ECG level, building on previous studies (Martinez-Navarro et al., 2019; Riebel et al., 2022; Wang et al., 2021; Zhou et al., 2022). The framework allows for personalisation of anatomical, structural and electrophysiological properties to clinical data. The biventricular anatomy depicted in Figure 4A was obtained from clinical MRI (Mincholé et al., 2019). Ventricular fibre orientation was incorporated using the rule-based method by (Doste et al., 2019) following anatomical observations from (Streeter et al., 1970). The anatomical description of the Purkinje network was incorporated using the algorithm by (Berg et al., 2023; Camps et al., 2023) to achieve consistency between simulated and clinical QRS complexes, as shown in Figure 4A. CV in fibre, sheet and normal directions were 65, (Taggart et al., 2000), 38 and 47 cm/s (see supplementary Figure S9). Propagation across the PMJs occurred with a delay of 2 ms in the anterograde direction, similar to experimental values of 3-7 ms (Joyner & Overholt, 1985), and 1 ms in the retrograde direction.

Electrical propagation was simulated with the monodomain equation in the GPU-enabled open source MonoAlg3D simulator (Sachetto Oliveira et al., 2018). Cellular membrane kinetics were represented by the ToR-ORd model for human ventricular cells (Tomek et al., 2019, 2020), the Trovato model for Purkinje cells (Trovato et al., 2020) and the Paci2020 model for hPSC-CMs (Paci et al., 2020). Electrophysiological heterogeneity across the ventricles was modelled including APD gradients of 35 ms transmurally based on (Boukens et al., 2015; Franz et al., 1987) with 70 % endocardial and 30 % epicardial cells and in the apicobasal direction of 10 ms by scaling G_Ks_ (supplementary Figures S9 and S10).

**Figure 4.**
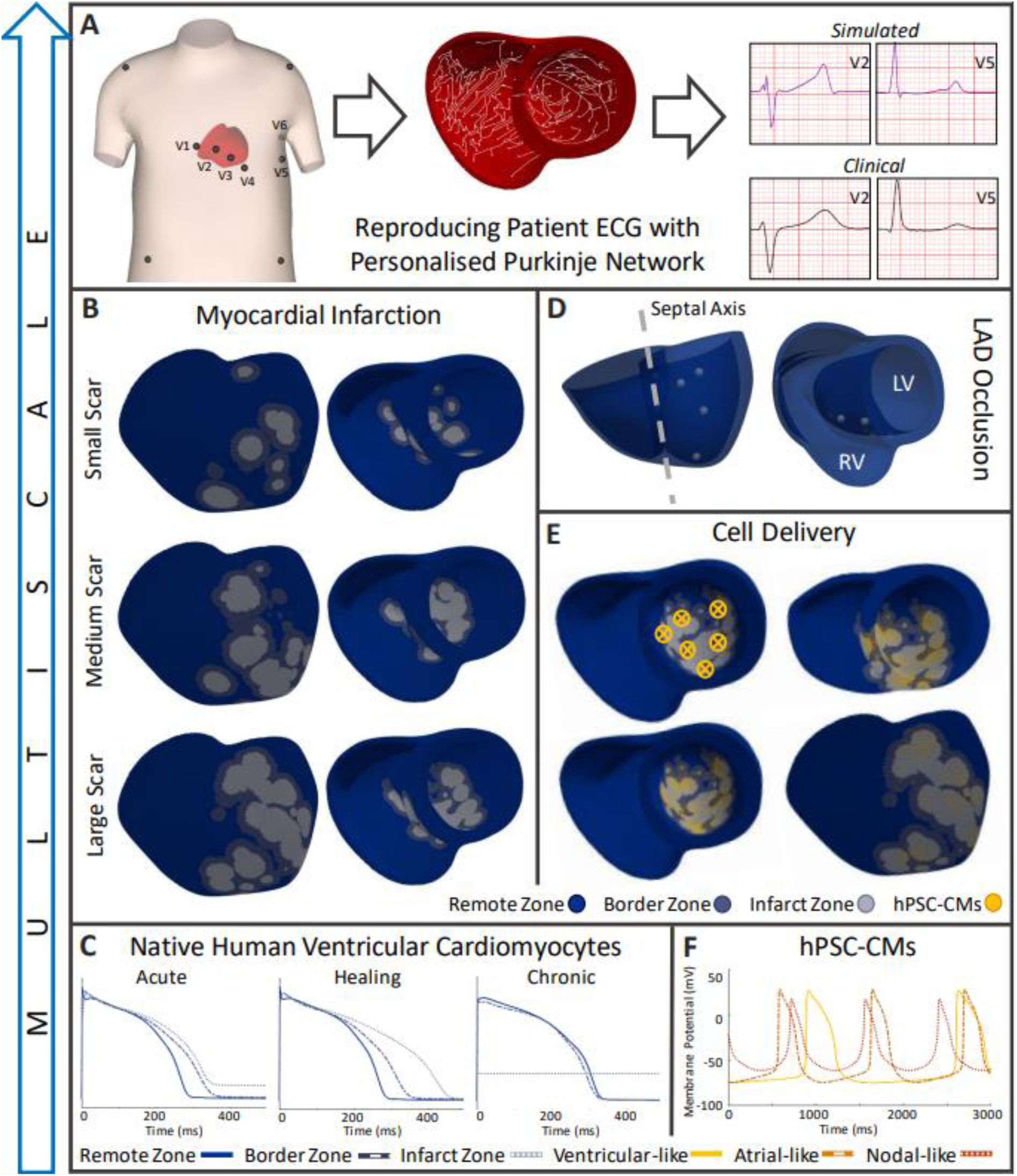
A) Patient heart and torso geometries with electrode locations (Mincholé et al., 2019) were combined with a personalised Purkinje conduction network to reproduce the patient’s ECG as in (Berg et al., 2023; Camps et al., 2023). B) Small, medium and large scar created using (Cardone-Noott et al., 2014; Hill et al., 2016). C) Action potential traces of the endocardial human adult ventricular ToR-ORd cell model during the acute, healing and chronic stages of myocardial infarction. D) Contact points representing the anatomy of the LAD; left (LV) and right (RV) ventricle indicated. E) Stem cell delivery locations in the left ventricle and delivered hPSC-CMs; basal, anterior and transmural view. F) Spontaneous action potential traces of the ventricular-like, atrial-like and nodal-like hPSC-CM Paci2020 model.

### Modelling stages of myocardial infarction

Three models of anteroseptal MI were produced corresponding to acute (first days), healing (first weeks) and chronic (several weeks) infarction (Figure 4C). Electrophysiological alterations were considered in the core of the infarct, the border zone and the remote myocardium (distant from the scar but affected over time). These were introduced by scaling conductances and time constants of key ionic currents, as reported experimentally and described in (Wang et al., 2021; Zhou et al., 2022), see Supplementary Table S1. The infarct zone during the chronic stage was modelled as non-electrically active but conducting (Rog-Zielinska et al., 2016) by setting the starting membrane potential to -60 mV (Ringenberg et al., 2014), accounting for the elevated DMP of fibroblasts which make up large proportions of myocardial scars (Kohl, 2003). Thus, the membrane potential in the chronic infarct changed only through diffusivity. Chronically infarcted elements produced AP-like changes in membrane voltage when coupled in tissue, which resembled experimentally and computationally observed behaviour of fibroblasts electrically coupled to cardiomyocytes (Klesen et al., 2018; Yue et al., 2011).

Using the algorithms described in (Cardone-Noott et al., 2014; Hill et al., 2016), three physiological scar geometries were produced and referred to as small, medium and large (Figure 4B). For this, six landmarks representing contact points of the left anterior descending artery (LAD) and the myocardium (Brinkman et al., 1994) were considered. As shown in Figure 4D, two contact points in the septum, three on the anterior wall and one in the apex were selected; apical involvement in anterior scars was highlighted in clinical studies such as (Ørn et al., 2007). To create the three scar sizes, the number of scar cores was set to 75, 200 and 275; further algorithm parameters are listed in supplementary Table S2. Scars covered 14, 23 and 27 % of the ventricles (composed of 5, 10 and 14 % infarct core and 9, 12 and 13 % border zone, respectively), in line with experimentally and clinically reported ranges (Reindl et al., 2020; Spath et al., 2021; Tülümen et al., 2021).

Diffusivity in the infarct and border zone was decreased to match a 70 % reduced CV in the healing stage, based on clinical observations (Aronis et al., 2020; Jamil-Copley et al., 2015). The calibrated diffusivities were maintained in the acute and chronic stages, resulting in about 75 and 90 % reduced CV in the infarct centre due to further cellular remodelling (see supplementary Table S3 for a full list of CVs).

### Modelling cell therapy

Virtual cell delivery was introduced at six injection locations across the infarct, as shown in Figure 4E, to cover all scar sizes. Existing adult ventricular cells (ToR-ORd model) were randomly replaced by the hPSC-CMs (Paci model), with probabilities depending on infarct versus border zone, transmurality and distance to the nearest delivery location (see supplementary Table S4 for further detail). Probabilities were calibrated using experimental insights from (Poch et al., 2022), where in non-human primate tissue slabs, more injected cells settled in the infarct core compared to the border zone. Informed by Poch et al.’s in vivo porcine models, in our simulations, 28.4, 22.1 and 23.4 % (different due to scar morphology), of the core infarcted area was replaced by hPSC-CMs in the small, medium and large scar, respectively. In the border zone, 9.3, 9.3 and 8.8 % of cells were replaced. Poch et al. also showed injected cells to migrate through the transmural wall; we incorporated this finding by modelling hPSC-CMs to be most likely to locate in the subendocardium, as shown in Figure 4E. Overall, with decreasing scar size, our algorithm resulted in similar proportions but lower absolute numbers of infarct being replaced with hPSC-CMs.

To investigate the effect of hPSC-CM heterogeneity, as experimentally observed in (Ban et al., 2017; Jiang et al., 2020), three hPSC-CM populations were considered for each scar size with varying degrees of heterogeneity: a homogeneous population consisting of ventricular-like hPSC-CMs only, a moderately heterogeneous population (made up of 80 % ventricular-like and 10 % atrial and nodal-like hPSC-CMs each) and a severely heterogeneous population (containing 50 % ventricular-like and 25 % nodal and atrial-like hPSC-CMs each). CV of hPSC-CMs was calibrated to 10 cm/s (Hansen et al., 2018).

Cell heterogeneity was reproduced by creating a population of 10,000 single cell hPSC-CM models using the Paci2020 model as baseline and scaling key ionic current conductances between 0 and 5. From this population, an atrial-like and a nodal-like cell type were selected by calibrating against data from (Ma et al., 2011). The AP traces, biomarkers (after 1000 spontaneous beats to reach steady state) and corresponding current conductance scaling factors of the three resulting models can be seen in Figure 4F and supplementary Tables S5 and S6, respectively.

Spontaneous beats in the human ventricles were defined as beats originating from the hPSC-CMs and depolarising both ventricles. The onset of spontaneous beats was quantified as the time when hPSC-CMs depolarised above -45 mV (75 % of the nodal-like, i.e., most depolarised, DMP). DMP in this work was defined as the most negative membrane potential between two upstrokes.

### Verification, validation and uncertainty quantification

Human modelling and simulation is a mature and well-established technology (Arevalo et al., 2013; Passini et al., 2017; Roney et al., 2020; Wang et al., 2021), shown to accurately replicate disease conditions and make precise predictions into underlying mechanisms and optimal treatment. To demonstrate the credibility of our framework, we considered verification, validation and uncertainty quantification principles (Musuamba et al., 2021; Viceconti et al., 2021).

Correct single cell model transfer across solvers is depicted in supplementary Figure S11. Verification of the monodomain model solver used in this work was ensured through benchmark tests (Sachetto Oliveira et al., 2018) and CV convergence analysis (supplementary Figures S12 and S13).

The single cell models used here were constructed using data obtained in human preparations, clinically supporting their biological relevance. Their credibility is further shown through extensive consistency with experimental data in control conditions and under drug block (Paci et al., 2020; Passini et al., 2017; Tomek et al., 2019; Trovato et al., 2022). The three biventricular models from acute to chronic infarction demonstrate agreement with experimental ionic remodelling (Wang et al., 2021; Zhou et al., 2022) and clinical ECG data (supplementary Figure S3). The Purkinje system was introduced using experimental data and personalisation of the activation sequences yielding ECG biomarkers consistent with the clinical ECG for the corresponding anatomy (Camps et al., 2023) (supplementary Figure S10).

Regarding parameter uncertainty, sensitivity analyses were conducted for single cell model parameters in their original publications and in (Paci et al., 2021; Riebel et al., 2021). The sensitivity of the biventricular model to disabled retrograde Purkinje propagation was investigated and as shown in supplementary Figure S14, ECG traces under sinus pacing were almost identical. To allow for accurate comparisons throughout our simulations, parameters were varied in a controlled manner to test specific hypotheses and only where necessary. Specifically, the biventricular geometry, activation pattern, occluded areas and hPSC-CM delivery locations were kept constant but could be personalised to individual patients in future studies. To maintain their spatial distribution, hPSC-CM heterogeneity scenarios were applied on top of the randomly created hPSC-CM locations.

### Simulation software and protocol

In total, we conducted 550 simulations for precise investigation of arrhythmia mechanisms in human hearts. These considered 27 scenarios, composed of three scar geometries across three infarct stages for three different hPSC-CM population heterogeneities. Initial conditions in the biventricular geometry were taken from the single cell models after reaching steady state (500 and 1000 paced beats at 1 Hz in the ToR-ORd and Trovato model, respectively, and 1000 spontaneous beats in the Paci model). Each biventricular scenario was paced for 3 beats under 1 Hz at the bundle of His, reaching consistency between beats. To quantify arrhythmic risk, we observed whether hPSC-CMs beat spontaneously after the third beat, followed by a fourth beat applied at varying intervals ranging from 1620 to 1880 ms after the last sinus beat. This fourth beat is representative of slow heart rates between 32 and 37 beats per minute (bpm). If a re-entrant wavefront occurred following the bradycardic beat, the simulations were observed for a further two seconds to evaluate arrhythmia sustainability. To investigate the arrhythmic role of Purkinje coupling, which may be affected by MI (Kienzle et al., 1987), we ran our simulation set-up once with and once without retrograde Purkinje propagation by disabling current flow from the tissue into the Purkinje network at PMJ sites.

We also studied the effect of heart rate on spontaneous hPSC-CM activity. In the single cell models, 1000 s spontaneous activity followed by 1000 paced beats (at 50 to 100 bpm) were simulated with the proceeding spontaneous beat analysed. Increasing pacing was explored also in the biventricular model with the medium scar and the most heterogeneous hPSC-CM population across all infarct stages.

Simulations were run using the finite volume method monodomain GPU model solver MonoAlg3D (Sachetto Oliveira et al., 2018), see configuration settings in supplementary Table S7. ECG signals were simulated with MonoAlg3D, based on the pseudobidomain approach from (Bishop & Plank, 2011). Through CV convergence analysis, the human geometry was refined to 300 μm resolution to give a CV error of 12 % (see supplementary Figure S12). With this setup, one *in silico* second took eight hours to simulate using the Piz Daint Platform at the CSCS Swiss National Supercomputing Centre.

## Acknowledgements

This project was funded by a BBSRC PhD iCASE (BB/V509395/1) and Russell Studentship Agreement with AstraZeneca (R67719/CN001) to L. L. R., an Engineering and Physical Sciences Research Council doctoral award to J. C., an Oxford-Bristol Myers Squibb Fellowship to X. Z. (R39207/CN063), a Wellcome Trust Senior Research Fellowship in Basic Biomedical Sciences to B. R. (214290/Z/18/Z), the Oxford BHF Centre of Research Excellence (RE/13/1/30181), an NC3Rs Infrastructure for Impact Award (NC/P001076/1), the EPSRC project CompBioMed X (EP/X019446/1), the CompBioMed project (European Commission Horizon 2020 research and innovation programme, grant agreements No. 675451 and No. 823712) and by the Brazilian Government via CAPES, CNPq, FAPEMIG, UFSJ and UFJF. The Paci2020 model was provided by Dr. Michelangelo Paci and the Computational Biophysics and Imaging Group (CBIG) at Tampere University Foundation. Simulations were enabled through PRACE-ICEI grants icp013 and 019 and carried out at the Piz Daint Platform of the Swiss National Supercomputing Centre.

This research was funded in part by the Wellcome Trust [grant number 214290/Z/18/Z]. For the purpose of open access, the author has applied a Creative Commons Attribution (CC BY) public copyright licence to any Author Accepted Manuscript version arising from this submission.

