## Supplementary Material for "*In Silico* Evaluation of Cell Therapy in Acute versus Chronic Infarction: Role of Automaticity, Heterogeneity and Purkinje in Human"

### Supplements

#### Supplementary Videos

Supplementary Video SV1 shows the spontaneous hPSC-CM beat after 3 sinus beats (starting at the bundle of His and with healthy Purkinje-myocyte coupling) in the large acute scar and most heterogeneous hPSC-CM population.

Supplementary Video SV2 depicts the spontaneous beat from video SV1 followed by a bradycardic sinus beat 1.81 s after the last sinus beat which re-enters through a PMJ.

#### Results - Simulations of progressing myocardial infarction

This section shows additional validation of post myocardial infarction through experimental ionic remodelling evidence, ECG and activation and repolarisation time maps.

***Table S1.*** *Scaling factors of ionic current conductances and time constants to model infarct, border and remote zones during the acute, healing and chronic stages, as established in* (Wang et al., 2021; Zhou et al., 2022). *Supporting experimental data can be found in* [1](Hund et al., 2008) [2](M. Jiang, 2000) [3](Maier et al., 2003) [4](Tomek et al., 2019) [5](Valdivia et al., 2005) [6](Hegyi et al., 2018) [7](M. T. Jiang et al., 2002) [8](Høydal et al., 2018) [9](Hoch et al., 1999) [10](Ringenberg et al., 2014).

| Scaled Parameter | Scaling Factor | | | | | | | | |
|---|---|---|---|---|---|---|---|---|---|
| | Acute | | | Healing | | | Chronic | | |
| | IZ | BZ | RZ | IZ | BZ | RZ | IZ | BZ | RZ |
| $V_m$ (set not scaled) | | | | | | | -60[10] | | |
| $K_o$ (set not scaled) | 8.5 | | | | | | | | |
| $G_{Na}$ | 0.575 | 0.575 | | 0.4[1] | 0.4[1] | | | 0.435[5] | 0.435[5] |
| $G_{NaL}$ | | | | | | | | 1.275[6] | 1.413[6] |
| $G_{to}$ | 0[1] | 0[1] | | 0[1] | 0[1] | | | | |
| $P_{Ca}$ | 0.7 | 0.7 | | 0.64[1] | 0.64[1] | | | 0.7[6] | |
| $G_{Kr}$ | 0.375[2] | 0.7 | | 0.375[2] | 0.7 | | | 0.89[6] | 0.87[6] |
| $G_{K1}$ | 0.6[1] | 0.6[1] | | 0.6[1] | 0.6[1] | | | 0.76[6] | |
| $J_{up}$ | | | | | | | | 0.4[7,8] | 0.4[7,8] |
| $G_{KCa}$ | | | | | | | | 2[6] | 2[6] |
| $G_{ClCa}$ | | | | | | | | 1.25[6] | 1.25[6] |
| aCaMK | 1.5[1] | 1.5[1] | | 1.5[1] | 1.5[1] | | | 1.5[9] | 1.5[9] |
| tau_relp | 6[3] | 6[3] | | 6[3] | 6[3] | | | 6[3] | 6[3] |
| $P_{Cab}$ | 1.33[4] | 1.33[4] | | 1.33[4] | 1.33[4] | | | | |

*IZ: infarct zone, BZ: border zone, RZ: remote zone*

As shown in Figures S1-S3, our simulations captured clinically observed T-wave inversion, fractured QRS-complexes, ST-segment elevation and QRS-complex prolongation. Our simulations further showed a notched T-wave in the acute and healing stages, where the clinical records (Figure S3B & C) showed a more biphasic than notched T-wave.

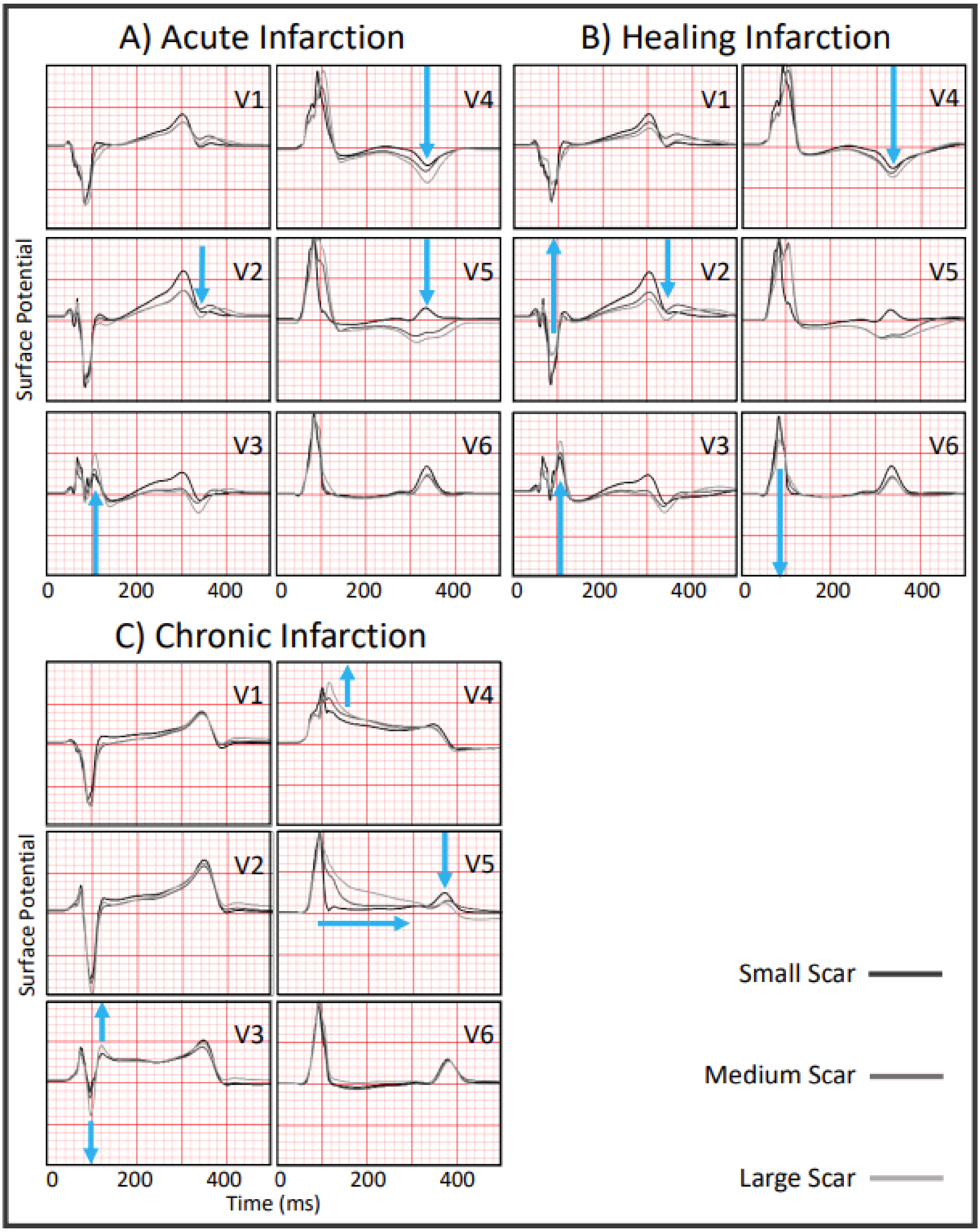

***Figure S1.*** *Comparison of ECG signals for the healthy control, small scar, medium scar and large scar across infarct stages with retrograde propagation enabled; blue arrows indicate the trend from the small to large scar ECG.*

Figure S2 below shows activation and repolarisation time maps as well as an ECG comparison from healthy across the different stages of infarction. Please note that activation and repolarisation times in grey were not calculated due to low voltage amplitude. While in the chronic stage current flowed into the infarct centre, the infarct cores did not depolarise above 0mV, due to the infarct being slowly conducting but electrically non-active (Figure S2C).

As the hPSC-CMs have a more depolarised diastolic membrane potential, we chose an activation threshold of -45 mV (75 % of the nodal-like, most depolarised, hPSC-CM phenotype). For comparability, we used the same voltage threshold to define activation and repolarisation across the scenarios before and after cell delivery, hence the repolarisation marks the repolarisation at 70 % in adult ventricular cardiomyocytes.

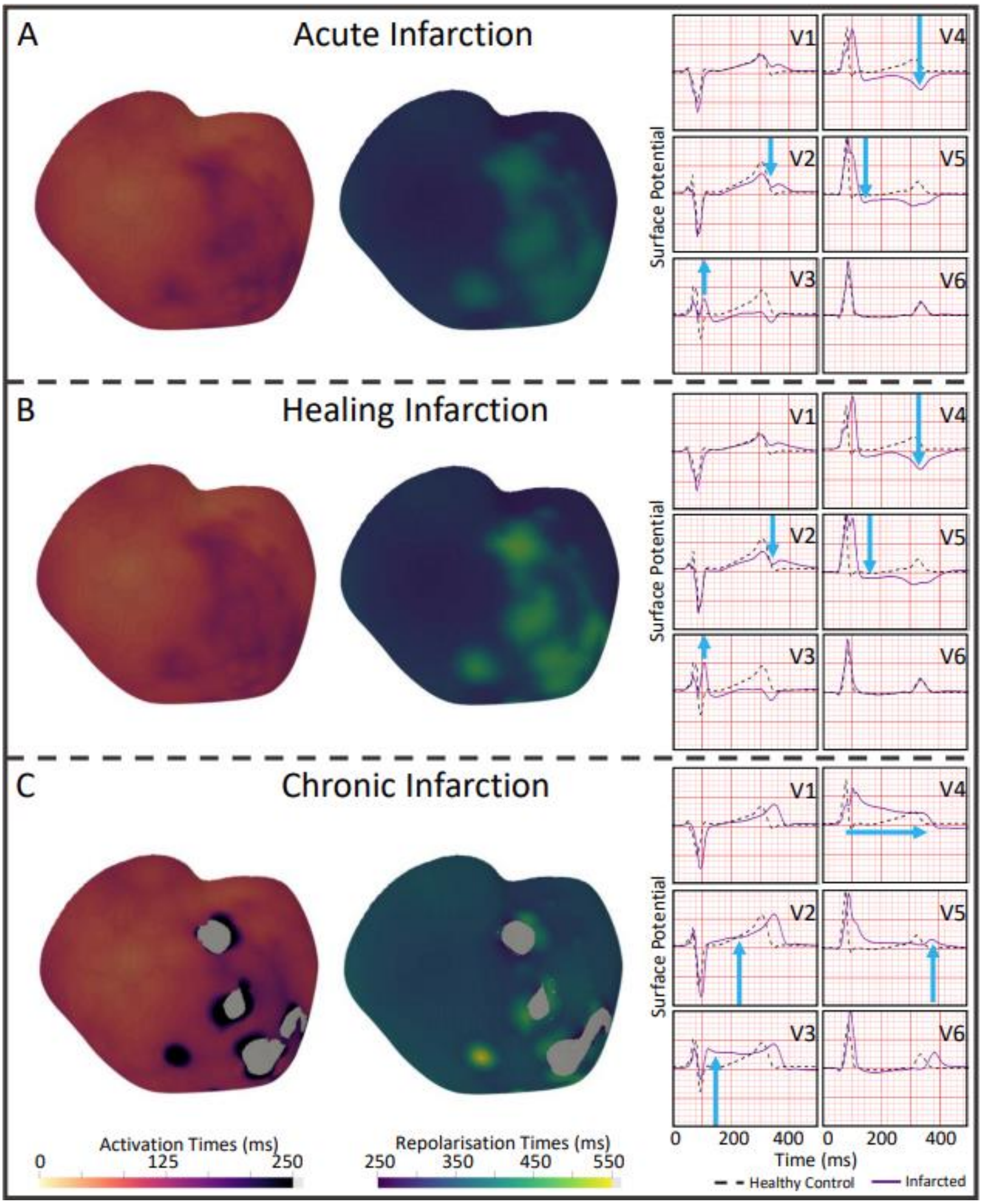

***Figure S2.*** *Activation and repolarisation time maps in the medium scar during the acute, healing and chronic stages of myocardial infarction with retrograde Purkinje propagation enabled and paced at 1Hz with the 3$^{rd}$ beat shown. Simulated ECG traces show the healthy control compared to the different infarct stages. An activation threshold of -45 mV was used to compute the activation time maps.*

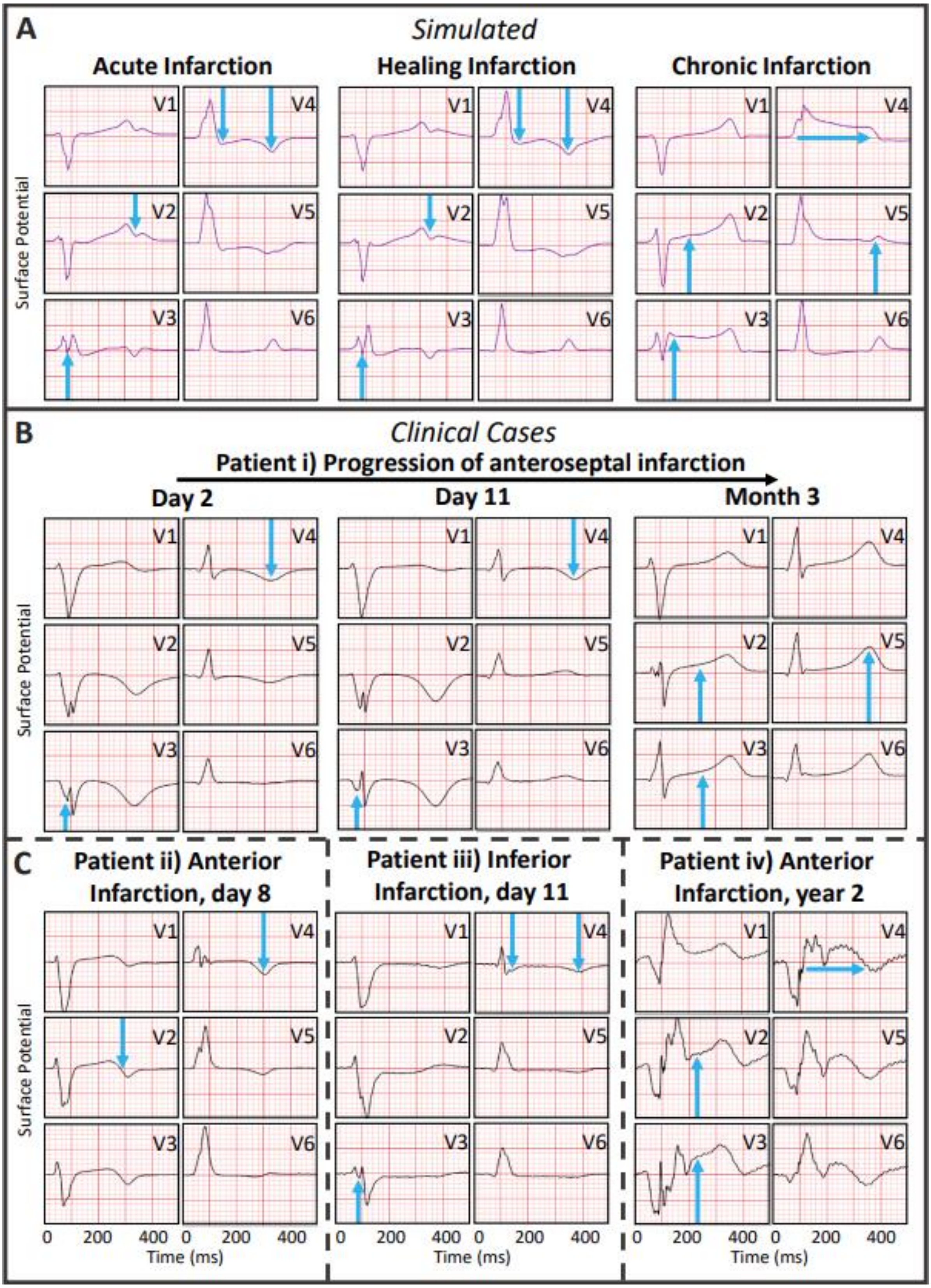

***Figure S3.*** *A) Simulated ECGs for the medium scar across infarct stages with retrograde Purkinje propagation enabled. B) Clinical ECG from the PTB Diagnostic ECG Database, https://physionet.org/content/ptbdb/1.0.0/* (Bousseljot et al., 2009; Goldberger et al., 2000)*, i) in patient 033 1, 10 and 89 days after antero-septal infarction. C) different patients: ii) Patient 163 7 days after anterior infarction, iii) patient 026 10 days after inferior infarction and iv) patient 148 21 months after anterior infarction. Note, A) and B) are the same as in Figure 1 but were included here again for a full comparison.*

#### Results - Simulations of arrhythmic risk after spontaneous hPSC-CM beats

This section provides further information on the effects of cell therapy on the simulations under control and arrhythmias.

The main difference between the different hPSC-CM phenotypes apart from their spontaneous beating rate, was their action potential duration. Hence, as shown in Figure S4 below, slight changes across the different hPSC-CM population heterogeneities were present in the repolarisation phase, i.e., the ECG's T-wave.

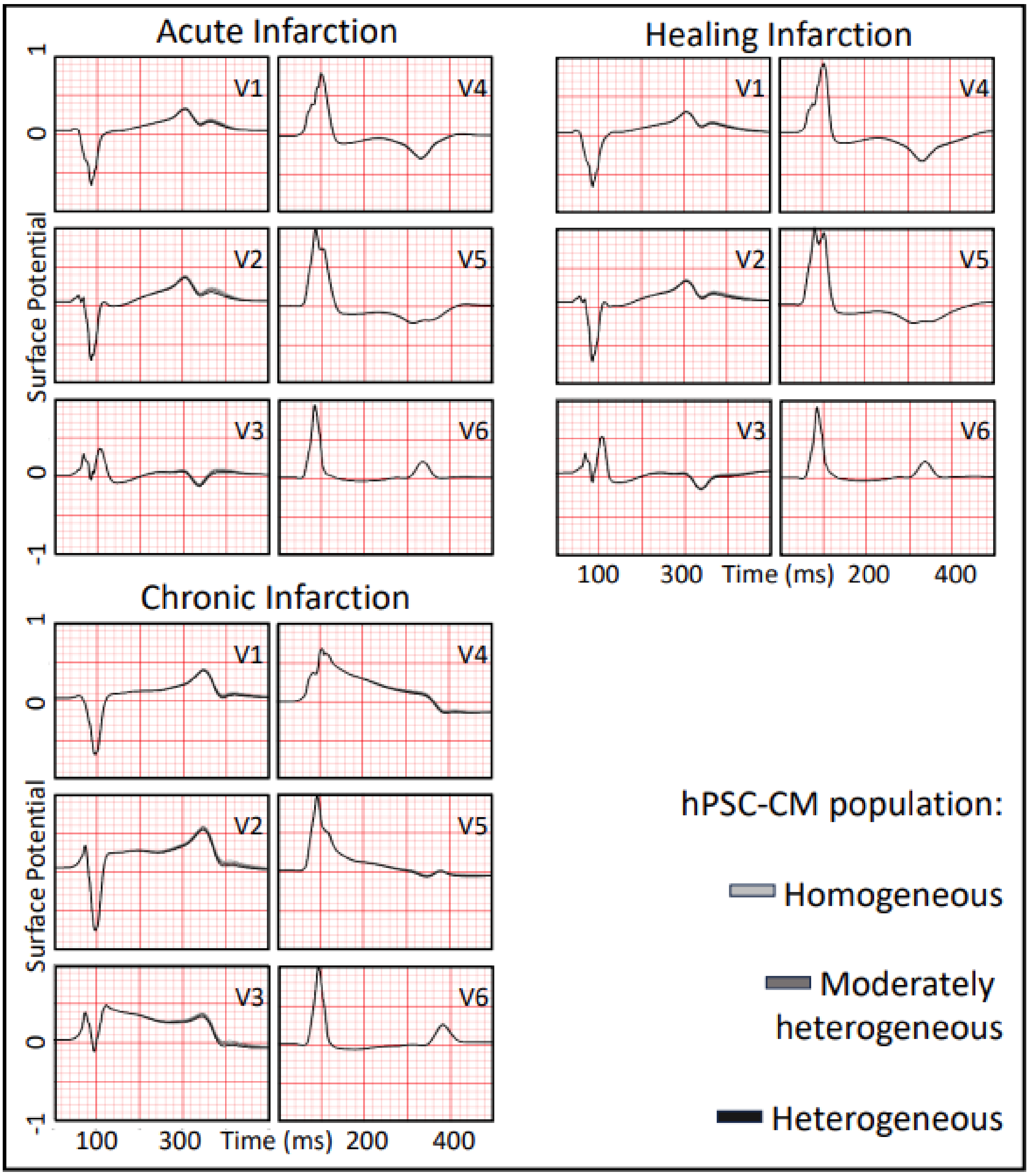

*Figure S4.* *Comparison of ECG signals in the medium scar across infarct stages after a homogenous (ventricular-like hPSC-CMs only, light grey traces), a moderately heterogeneous (80 % ventricular-like, 10 % atrial-like and 10 % nodal-like hPSC-CMs, dark grey traces), or a severely heterogeneous (50 % ventricular-like, 25 % atrial-like and 25 % nodal-like hPSC-CMs, black traces) hPSC-CM population was introduced.*

As shown in Figure 2B and the activation time maps in supplementary Figure S5 below, while most hPSC-CM delivery sites depolarised spontaneously, the septal site was the first to do so for all infarct stages and scar sizes. This is due to the septum being activated first during sinus rhythm originating at the bundle of His. A large absolute number of hPSC-CM in the septal wall due to its thickness further favoured spontaneous depolarisation. Figure S5 below also highlights that while hPSC-CMs in the chronic stage depolarise spontaneously first (bright colours), propagation in the acute and healing stages was faster (more purple colour distribution) in the infarct as well as the remote zone.

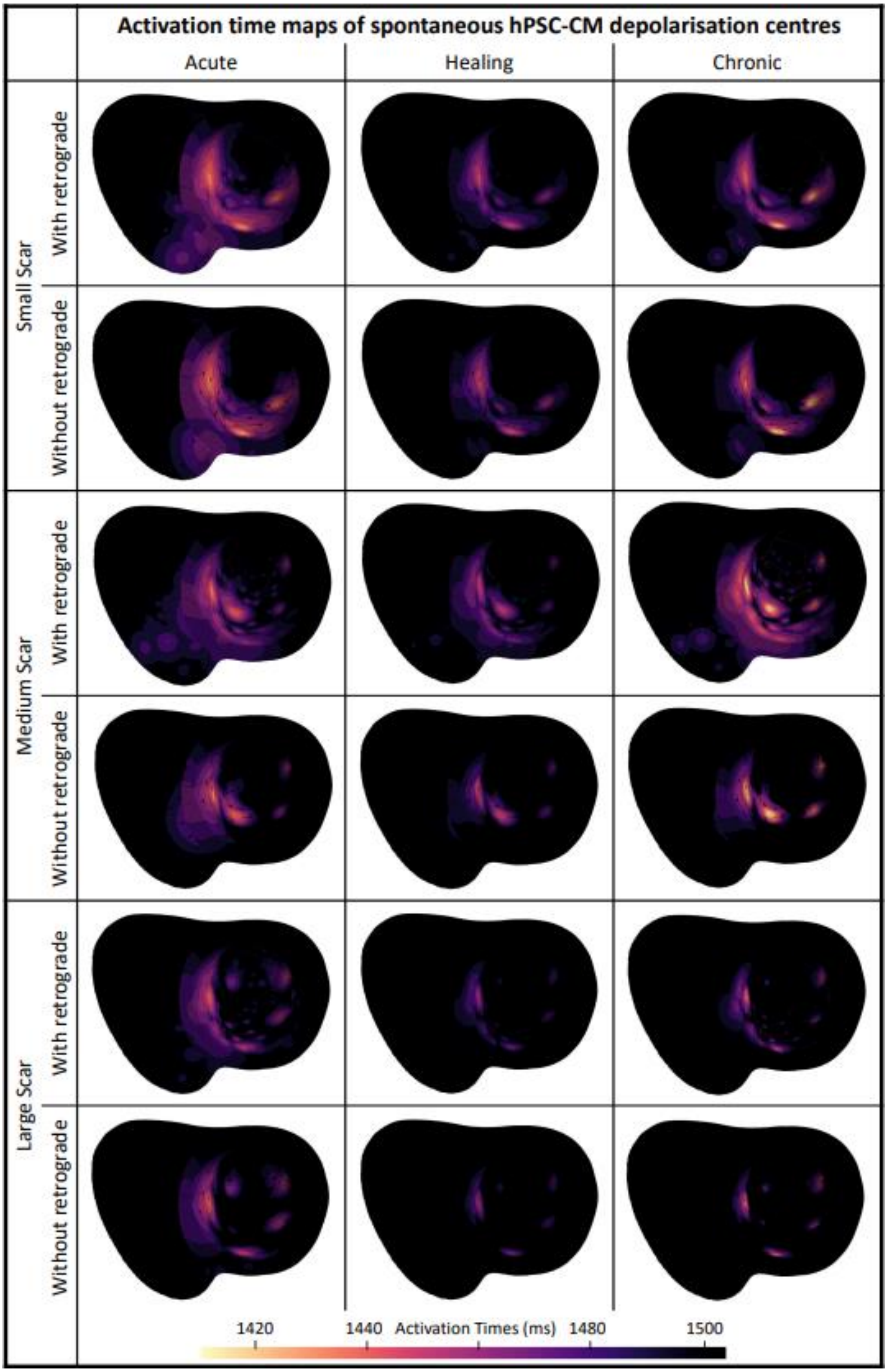

***Figure S5.*** *Activation time maps (following the last regular sinus beat) of spontaneous hPSC-CM depolarisation locations across infarct stages and scar sizes with and without retrograde Purkinje propagation enabled; activation times over 1500 ms are shown in black. Activation was determined through a -45 mV threshold.*

As shown in the text in Figure 2A, infarct size did not affect the occurrence of spontaneous beats, but their rate, which was slowest in the large scar. In Figure S6 below we show that this is due to the infarct's repolarisation times, determined by activation sequence (and thus CV) and action potential duration (APD). In chronic MI, lack of electrical activity in the infarct and reduced fast sodium current in remote myocardium caused delayed activation as well as APD prolongation, compared to acute and healing infarction. APD was also prolonged in the infarct centre during healing MI because of remodelling of potassium channels. In larger scars, the infarct provided a stronger current source against the surrounding healthy tissue sink, strengthening such APD differences. Due to electrotonic interactions, this resulted in mean hPSC-CM repolarisation times being shortest in the small scar during the acute stage, 375.51 ± 14.06 ms, and longest in the large scar during the chronic stage, 497.98 ± 76.13 ms.

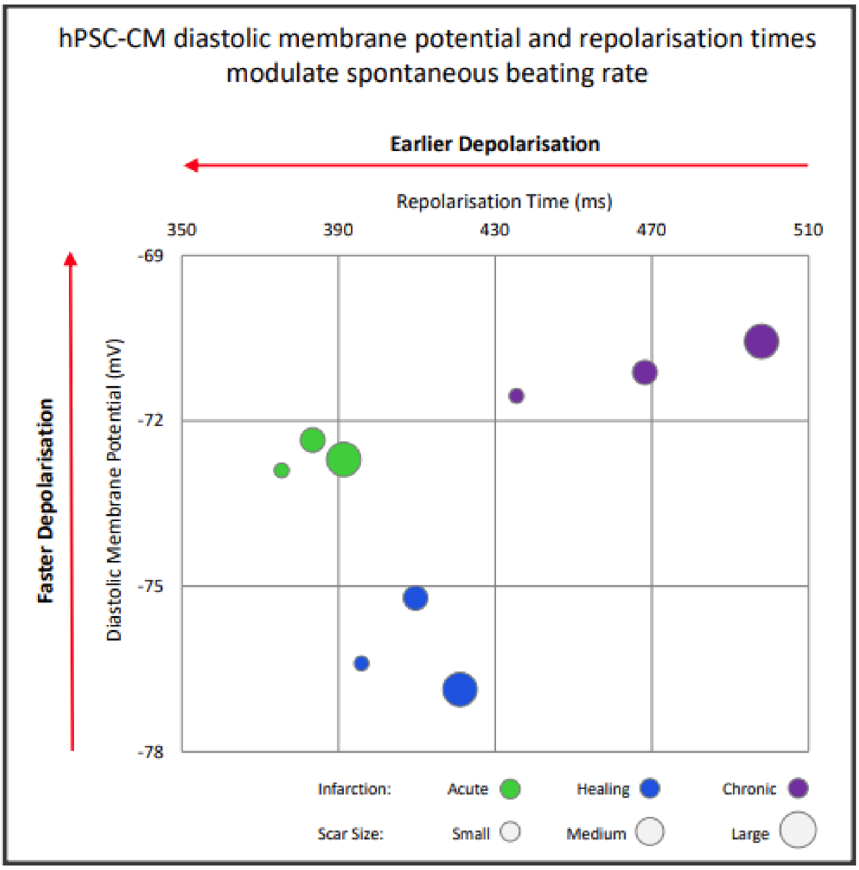

*__Figure S6.__ Two main mechanisms explain shortening of hPSC-CM beating intervals across infarct stages and sizes: average repolarisation time following sinus pacing versus most depolarised diastolic membrane potential of hPSC-CMs in the infarct centre after the last sinus beat. Red arrows indicate the direction of increased spontaneous beating.*

Figure S7 shows a comparison of ionic currents modulating the spontaneous beating rate following different pacing rates in the ventricular-like hPSC-CM phenotype. As identified in previous studies (Kim et al., 2015; Paci et al., 2018), hPSC-CMs spontaneously depolarise due to the funny current ($I_f$) as well as the cells' intrinsic calcium dynamics. Our simulations show that similarly to adult cardiomyocytes (Vassalle & Lin, 2004) faster pacing rate causes an intracellular calcium accumulation, as shown in the inlay in Figure S7 top right panel. Due to the faster pacing, the calcium uptake ($I_{up}$) in the sarcoplasmic reticulum (SR) was reduced and less calcium released from the SR through $I_{rel}$, see middle row left and middle panel. This caused a lower intracellular calcium concentration peak, as shown in the top row right panel. The sodium-calcium exchanger ($I_{NaCa}$) and sodium currents' ($I_{Na}$ and $I_{NaL}$) peak were also reduced, as shown in the bottom row panels. These changes in calcium dynamics explain the subsequent delayed spontaneous depolarisation.

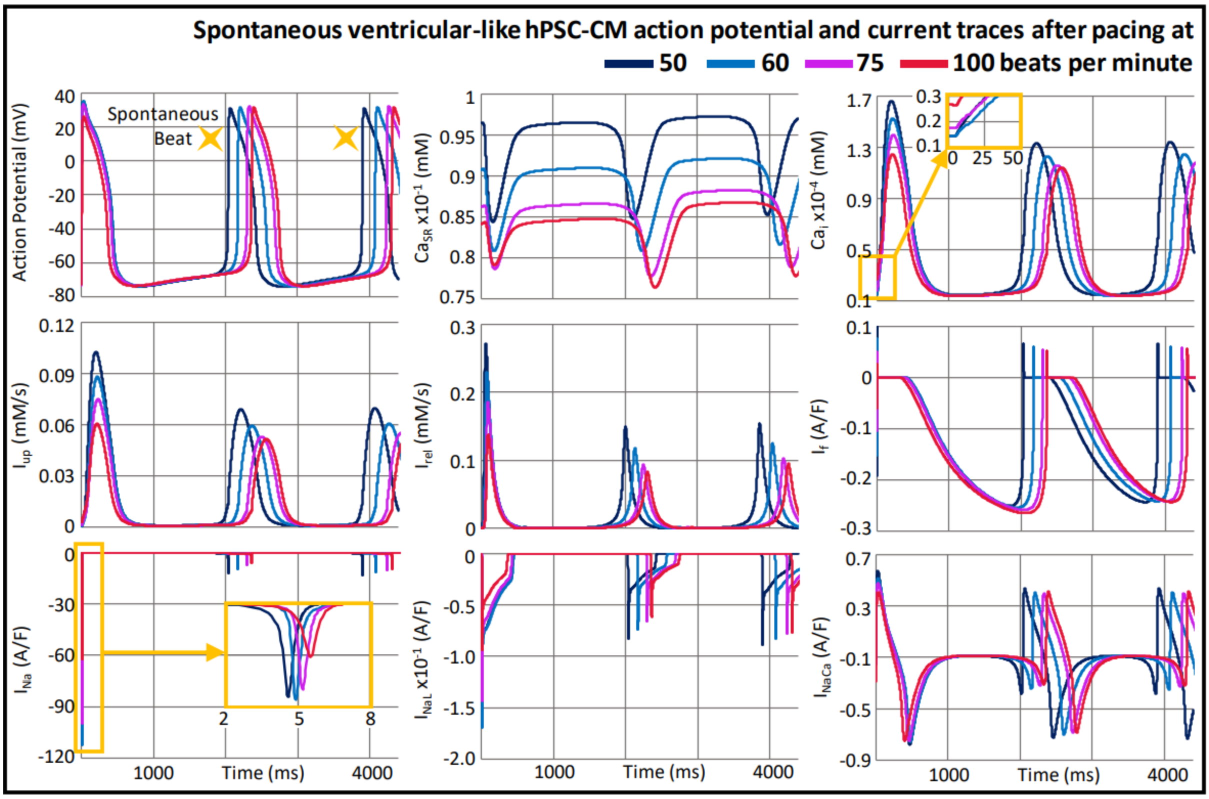

*Figure S7.* *Action potential and ionic current traces for the ventricular-like hPSC-CM phenotype; shown is the last of 1000 sinus beats at 50 (dark blue), 60 (light blue), 75 (purple) or 100 (red) bpm pacing followed by two spontaneous beats.*

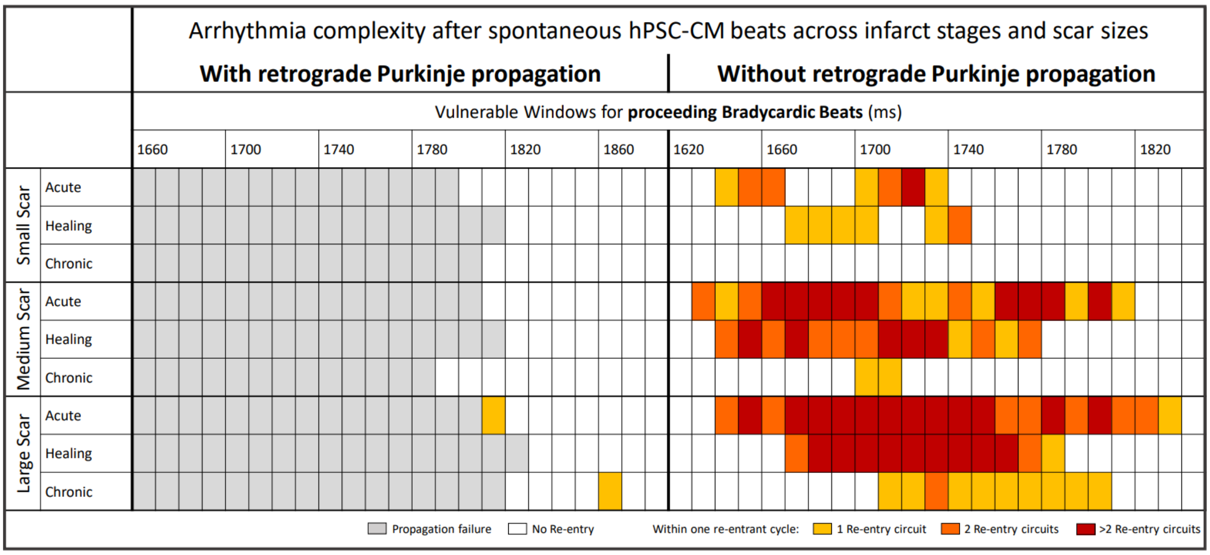

***Figure S8.*** *Arrhythmia complexity across all infarct stages and scar sizes and with and without retrograde Purkinje propagation enabled; measured as the number of re-entrant circuit locations within the first re-entrant cycle.*

#### Methods - Multiscale modelling and simulation of human electrophysiology

This supplementary section illustrates details of the electrophysiological human modelling and simulation framework under control conditions.

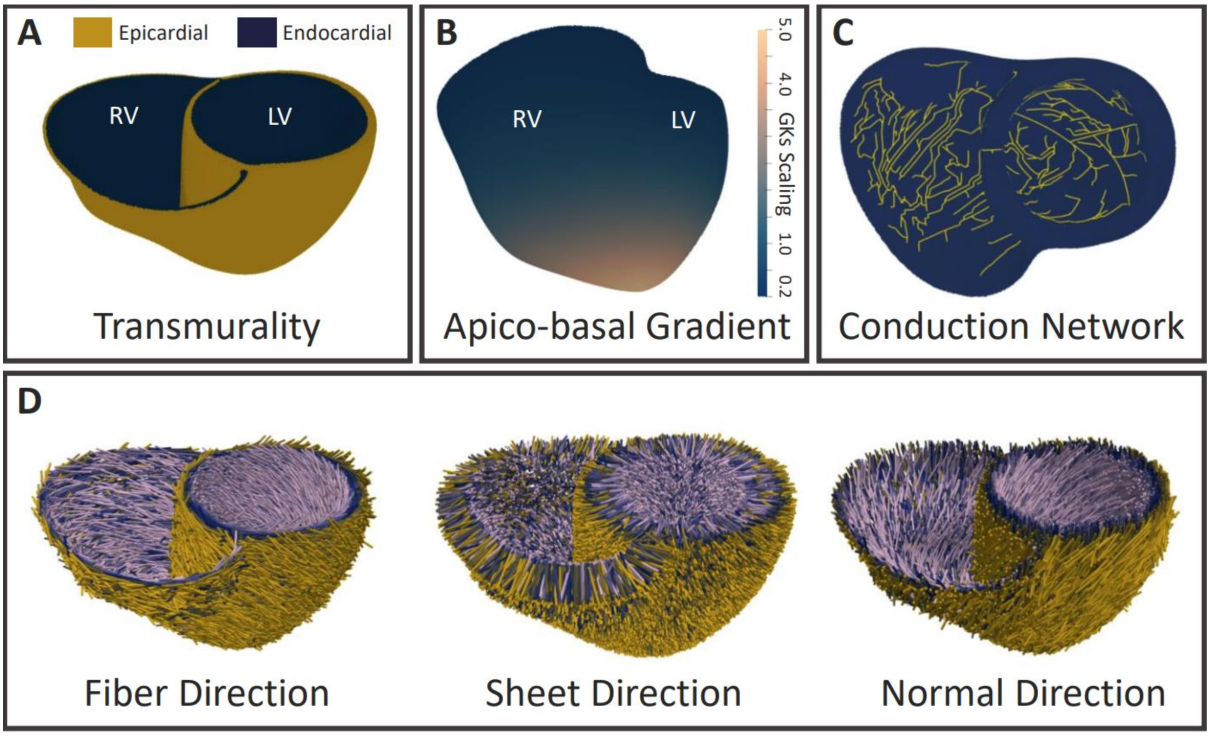

***Figure S9.*** *A) Transmural composition of the heart wall; 70% endocardial and 30% epicardial cells; left (LV) and right (RV) ventricle indicated. B) Scaling of the slow delayed rectifier potassium current, $I_{Ks}$, to reproduce an apico-basal action potential duration gradient. C) In silico cardiac conduction network* (Berg et al., 2023; Camps et al., 2023). *D) Fibre, Sheet and Sheet Normal direction of cardiac muscle fibres* (Doste et al., 2019).

Note in Figure S10 below, as the hPSC-CMs have a more depolarised diastolic membrane potential, we chose an activation threshold of -45 mV (75 % of the nodal-like, most depolarised, hPSC-CM phenotype). For comparability, we used the same voltage threshold to define activation and repolarisation across the scenarios before and after infarction and cell delivery, hence the action potential duration is that at 70 % repolarisation in adult ventricular cardiomyocytes.

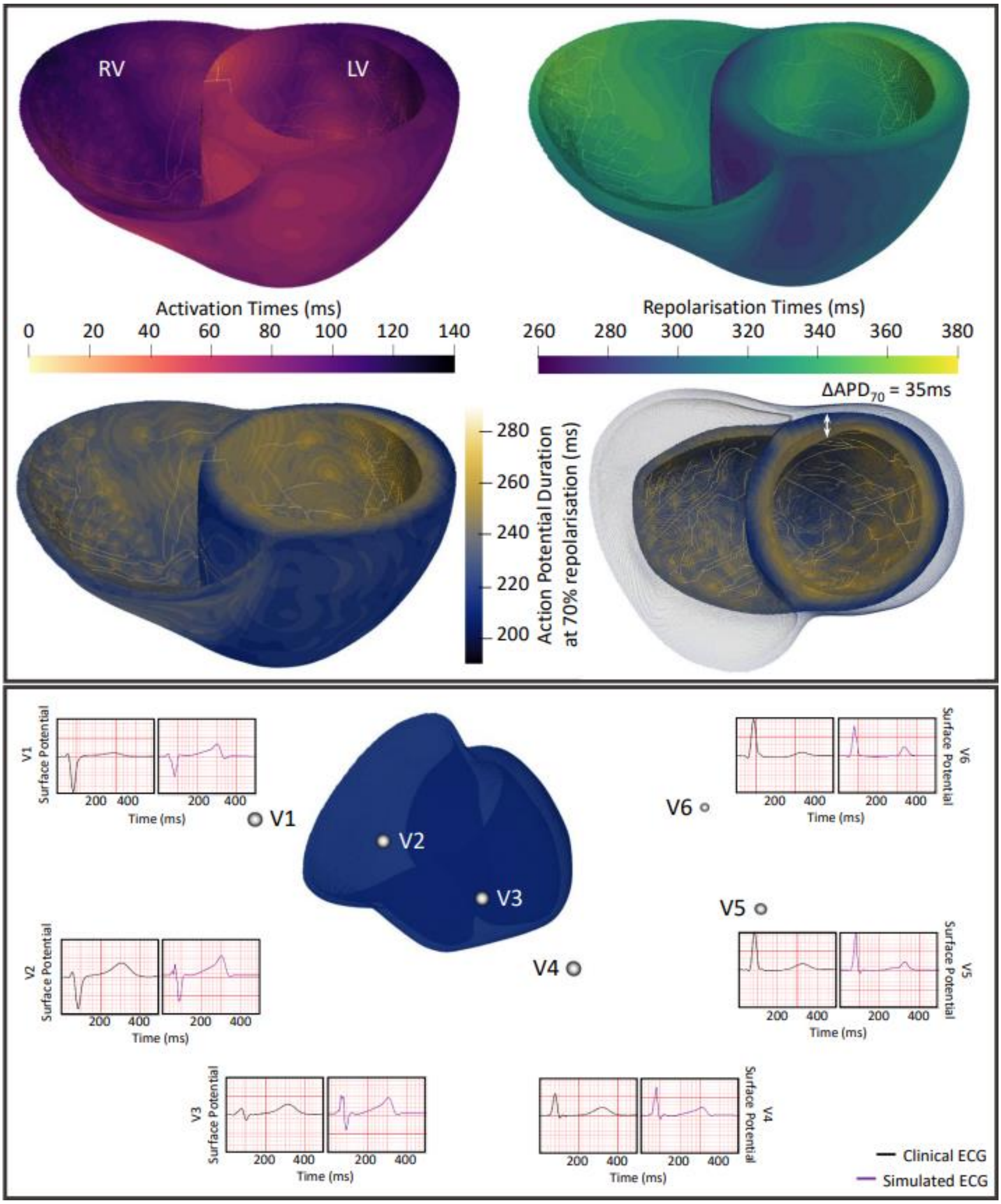

***Figure S10.*** *Top: Activation and repolarisation time maps as well as action potential durations at 70% repolarisation in the healthy control biventricular simulations. To determine activation, a threshold of -45 mV was used (this*

*maintained consistency with later activation thresholds for hPSC-CMs). $\Delta$$APD_{70}$ refers to the difference between endo and epicardial $APD_{70}$; left (LV) and right (RV) ventricle indicated. Bottom: Electrode positions as well as clinical and simulated ECG traces in the healthy control scenario.*

#### Methods - Modelling and simulating myocardial infarction

This section includes information on how different scars were created using the algorithm by (Cardone-Noott et al., 2014; Hill et al., 2016).

***Table S2.*** *Scar algorithm* (Cardone-Noott et al., 2014; Hill et al., 2016) *parameter values to create the small, medium and large scar.*

| Parameter | Small Scar | Medium Scar | Large Scar |
|---|---|---|---|
| Core_ext_noise | | 6 | |
| Core_post_noise | | 0 | |
| CZ_threshold | | 0.13 | |
| BZ_threshold | | 0.18 | |
| ncores | 75 | 200 | 275 |

***Table S3.*** *Conduction velocity (cm/s) for each infarct stage and zone (all considering the endocardial cell type and calibrated in a tissue slab). Remote zone diffusivity tensor: (2.60, 1.05, 1.60) mS/cm, infarct and border zone diffusivity tensor: (0.60, 0.35, 0.45) mS/cm, stem cell diffusivity tensor: (0.2, 0.2, 0.2) mS/cm. F: fibre, S: sheet (transmural), N: sheet normal direction.*

| | Conduction Velocities (cm/s) | | | | | | | | |
|---|---|---|---|---|---|---|---|---|---|
| | Remote Zone | | | Border Zone | | | Infarct Zone | | |
| | F | S | N | F | S | N | F | S | N |
| Acute | 65 | 38 | 47 | 22 | 14 | 17 | 15 | 10 | 12 |
| Healing | 65 | 38 | 47 | 19 | 12 | 15 | 19 | 12 | 15 |
| Chronic | 49 | 27 | 34 | 19 | 12 | 15 | 9 | 7 | 8 |
| | F | | | S | | | N | | |
| hPSC-CMs | 10 | | | 10 | | | 10 | | |

#### Methods - Modelling and simulating cell therapy

This section outlines how cell therapy was simulated at organ level and how different single cell stem cell phenotypes were created. To model cell delivery, a uniformly distributed random number was drawn for each mesh element, using Matlab's rand function. The element was declared an hPSC-CM, if this random number was smaller than the probability to be assigned the hPSC-CM model (based on the distance to the nearest injection location, transmurality and infarct location, as detailed in supplementary Table S4 below). Probabilities were calibrated to achieve a transmural and infarct to border zone ratio using experimental data from (Poch et al., 2022). In our algorithm, no hPSC-CMs were modelled in the right ventricle, considering the entire septum as part of the left ventricle. Next, the algorithm selected which hPSC-CM phenotype should be used, based on the hPSC-CM population-dependent percentages. I.e., for the most heterogeneous population, the probability was 50, 25 and 25 % to be assigned the ventricular-like, atrial-like and nodal-like phenotype model, respectively.

***Table S4.*** *Stem cell delivery algorithm parameter values.*

| | | | |
|---|---|---|---|
| **Assigned hPSC-CM if:** | Random number between 0 and 100 < distance-dependent scaling factor * area-dependent probability * 100 | | |
| **Distance-dependent scaling factor** | $3 - \frac{\text{minimal distance to any injection location}}{5500}$ | | |
| **Area-dependent probability** | | | |
| | Infarct Zone | Border Zone | Remote Zone |
| Transmurality (30:40:30) | | | |
| Endocardial | 0.21 | 0.09 | 0.00 |
| Subendocardial | 0.36 | 0.16 | 0.00 |
| Epicardial | 0.11 | 0.04 | 0.00 |

To incorporate different phenotypes of hPSC-CMs, a population of hPSC-CM Paci2020 models was created, by scaling key current conductances between 0.0 and 5.0. Specifically, the scaled conductances were those of the fast and late sodium currents ($I_{Na}$ and $I_{NaL}$), the outward transient potassium current ($I_{to}$), the rapid and slow delayed outward potassium currents ($I_{Kr}$ and $I_{Ks}$), the inward rectifier potassium current ($I_{K1}$), the L-type calcium current ($I_{CaL}$), the sodium-calcium exchanger current ($I_{NaCa}$), the sodium-potassium pump current ($I_{NaK}$), the calcium uptake flux via the SERCA pump ($J_{up}$), the funny current ($I_f$), the calcium release from the sarcoplasmic reticulum flux ($J_{rel}$) and the sarcoplasmic reticulum calcium leak flux ($J_{leak}$). In addition to the baseline ventricular-like Paci2020 model, an atrial-like and nodal-like phenotype were selected from the population. To allow for comparability, we chose to calibrate those two new models against the dataset to which the baseline ventricular-like Paci2020 model was calibrated, which includes data also on atrial-like and nodal-like phenotypes as shown in supplementary Table S5. Additionally, the maximum spontaneous action potential upstroke velocity (dV/dt_max, not used for calibration) was 20.55, 8.92 and 4.30 mV/ms in the ventricular-like, atrial-like and nodal-like models, respectively. The ionic current conductance scaling factors can be seen in supplementary Table S6.

***Table S5.*** *Experimental (Ma et al., 2011) and simulated spontaneous action potential biomarkers of the ventricular-like, atrial-like and nodal-like induced pluripotent stem cell-derived cardiomyocytes.*

| | DMP (mV) | | Peak Voltage (mV) | | $APD_{90}$ (ms) | | Interval (s) | |
|---|---|---|---|---|---|---|---|---|
| | Exp. | Sim. | Exp. | Sim. | Exp. | Sim. | Exp. | Sim. |
| Ventricular-like | -76 | -75 | 28 | 27 | 415 | 402 | 1.7 | 1.71 |
| Atrial-like | -74 | -74 | 27 | 28 | 286 | 269 | 1.2 | 1.05 |
| Nodal-like | -60 | -61 | 19 | 17 | 255 | 271 | 1.0 | 0.84 |

*DMP: diastolic membrane potential, $APD_{90}$: action potential duration at 90 % repolarisation, Interval: spontaneous beating interval*

***Table S6.*** *Ionic current scaling factors to produce the ventricular-like, atrial-like and nodal-like phenotypes of the stem cell-derived cardiomyocyte Paci2020 model.*

| Scaled Parameters | Scaling Factor | | |
|---|---|---|---|
| | Ventricular-like Model | Atrial-like Model | Nodal-like Model |
| $G_{NaF}$ | 1.00 | 0.57 | 1.17 |
| $G_{NaLmax}$ | 1.00 | 3.48 | 1.84 |
| $G_{to}$ | 1.00 | 3.15 | 2.28 |
| $G_{Kr}$ | 1.00 | 4.21 | 3.15 |
| $G_{Ks}$ | 1.00 | 0.63 | 0.57 |
| $G_{K1}$ | 1.00 | 0.35 | 0.01 |
| $G_{CaL}$ | 1.00 | 1.75 | 0.95 |
| $k_{NaCa}$ | 1.00 | 1.54 | 1.18 |
| $P_{NaK}$ | 1.00 | 1.17 | 0.95 |
| $V_{maxup}$ | 1.00 | 2.96 | 0.05 |
| $G_{f}$ | 1.00 | 1.17 | 1.25 |
| $G_{irelmax}$ | 1.00 | 0.09 | 1.20 |
| $V_{leak}$ | 1.00 | 0.11 | 2.91 |

#### Methods - Simulation software, verification and validation

This section focuses on simulation software settings and provides additional supporting information on the framework verification and validation.

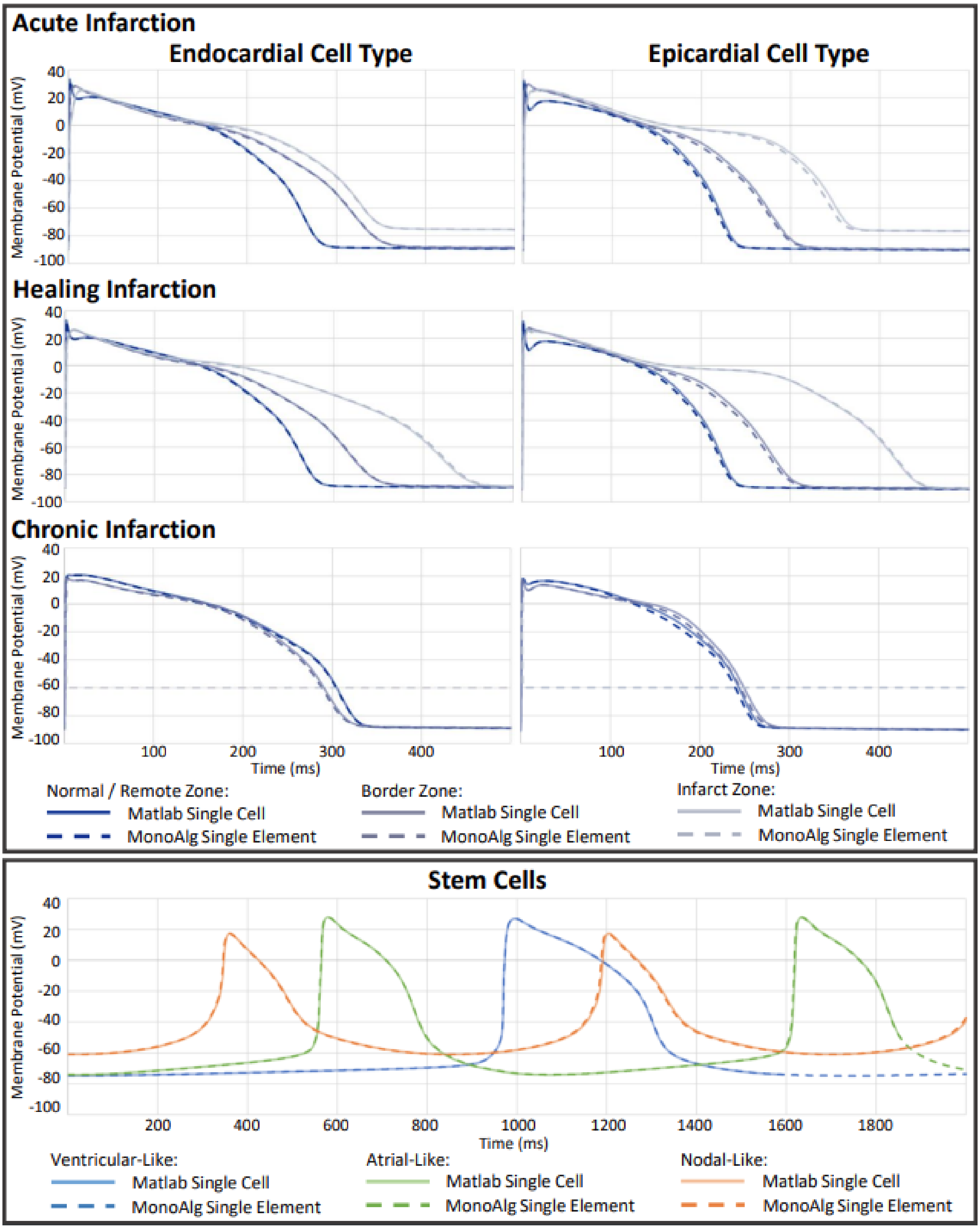

***Figure S11.*** *Action potential traces across infarction stages and cell types from the human ventricular ToR-ORd model and the stem cell-derived cardiomyocyte Paci2020 model solved in Matlab and the monodomain solver MonoAlg3D.*

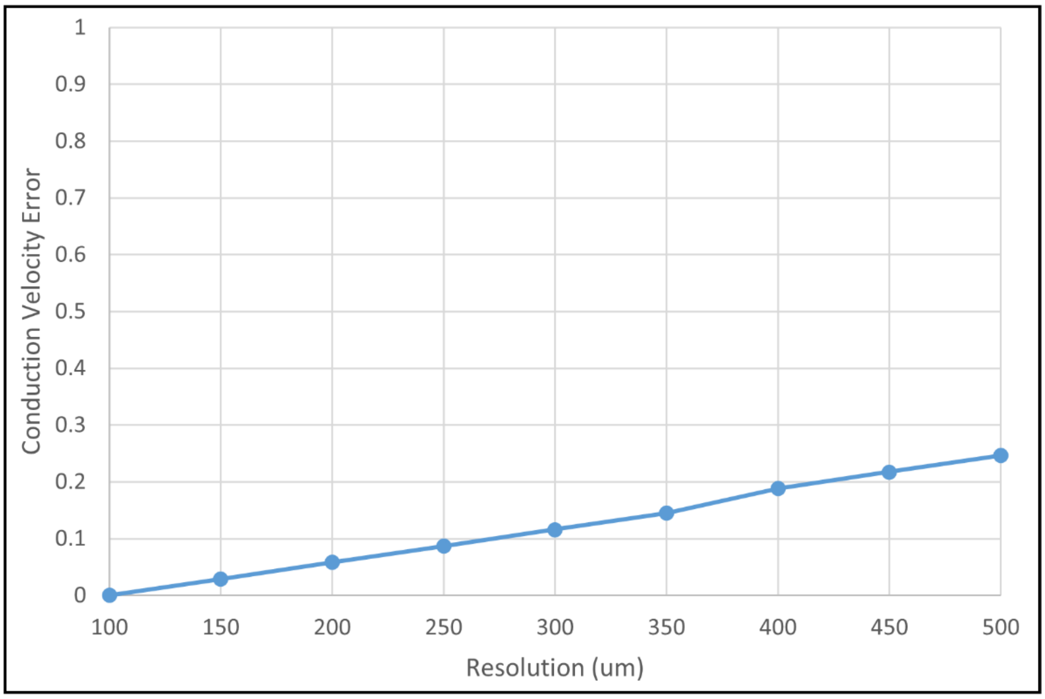

***Figure S12.*** *Conduction velocity error for the longitudinal conduction velocity of 70 cm/s across mesh resolutions in a 20 x 7 x 3 mm tissue slab. A 2 ms stimulus was applied in a corner of 1.5 x 1.5 x 1.5 mm.*

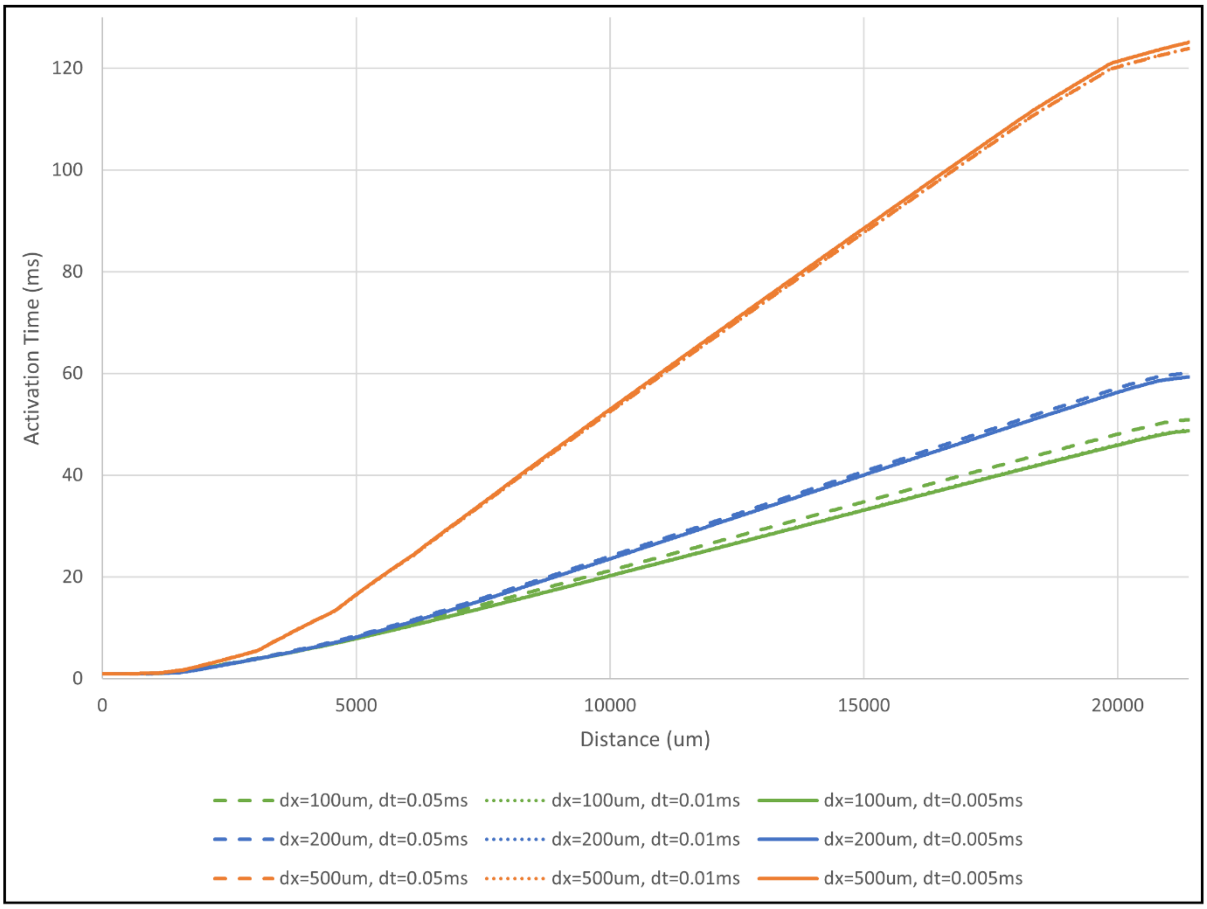

***Figure S13.*** *Benchmark analysis as in* (Niederer et al., 2011), *for varying mesh resolutions and time steps. A tissue slab with dimensions 20 x 7 x 3 mm was used and a 2 ms stimulus was applied in a 1.5 x 1.5 x 1.5 mm corner. Conductivities were set to transversely isotrop with $\sigma_l$ = 1.334 cm/s and $\sigma_t$ = $\sigma_n$ = 0.176 cm/s.*

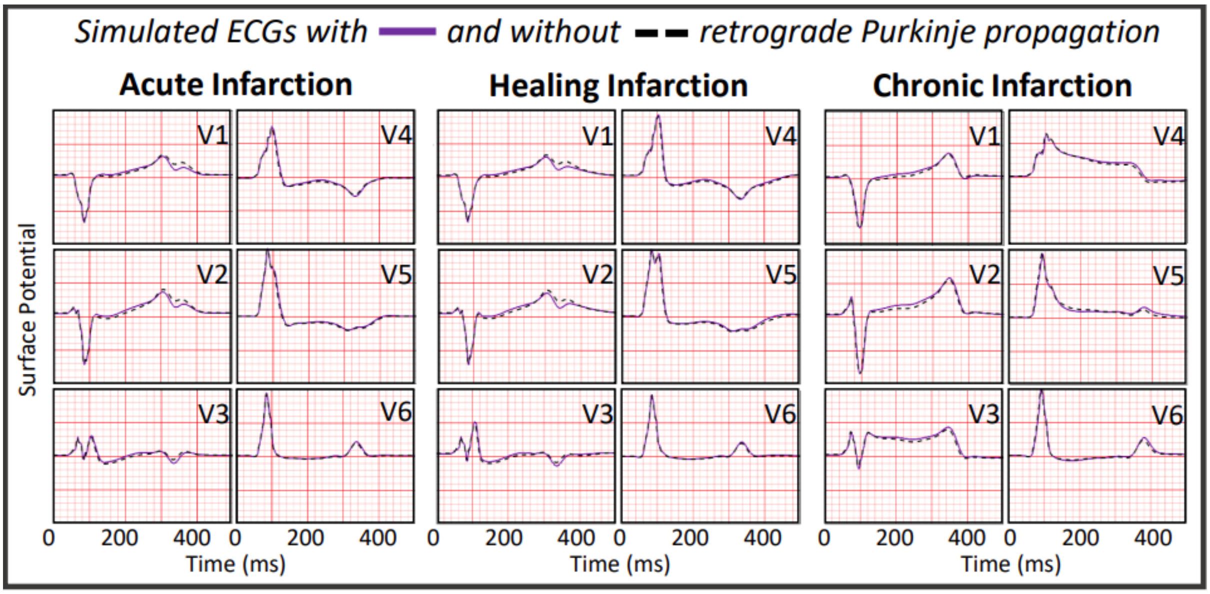

***Figure S14.*** *Comparison of simulated ECGs in the medium scar with the heterogeneous hPSC-CM population across all infarct stages with (solid purple line) and without retrograde Purkinje propagation (dotted black line) enabled.*

***Table S7.*** *MonoAlg3D configuration parameters for the biventricular simulations.*

| Parameter | ToR-ORd | Paci2020 | Trovato |
|---|---|---|---|
| ODE time step | 0.01 | 0.01 | 0.01 |
| ODE absolute tolerance | 1e-12 | 1e-12 | default |
| ODE relative tolerance | default | default | default |
| ODE solver method | Rush-Larsen + Euler | Rush-Larsen + Euler | Rush-Larsen + Euler |
| ODE adaptivity | false | false | false |
| Linear system solver function | biconjugate gradient | biconjugate gradient | conjugate gradient |
| Linear system solver tolerance | 1e-12 | 1e-12 | 1e-16 |
| PDE timestep | 0.01 | | |
| Space adaptivity | false | | |
| ODE solved in: | GPU | GPU | CPU |
| PDE solved in: | GPU | GPU | GPU |
